# Model-based genotype and ancestry estimation for potential hybrids with mixed-ploidy

**DOI:** 10.1101/2020.07.31.231514

**Authors:** Vivaswat Shastry, Paula E. Adams, Dorothea Lindtke, Elizabeth G. Mandeville, Thomas L. Parchman, Zachariah Gompert, C. Alex Buerkle

## Abstract

Non-random mating among individuals can lead to spatial clustering of genetically similar individuals and population stratification. This deviation from panmixia is commonly observed in natural populations. Consequently, individuals can have parentage in single populations or involving hybridization between differentiated populations. Accounting for this mixture and structure is important when mapping the genetics of traits and learning about the formative evolutionary processes that shape genetic variation among individuals and populations. Stratified genetic relatedness among individuals is commonly quantified using estimates of ancestry that are derived from a statistical model. Development of these models for polyploid and mixed-ploidy individuals and populations has lagged behind those for diploids. Here, we extend and test a hierarchical Bayesian model, called entropy, which can use low-depth sequence data to estimate genotype and ancestry parameters in autopolyploid and mixed-ploidy individuals (including sex chromosomes and autosomes within individuals). Our analysis of simulated data illustrated the trade-off between sequencing depth and genome coverage and found lower error associated with low depth sequencing across a larger fraction of the genome than with high depth sequencing across a smaller fraction of the genome. The model has high accuracy and sensitivity as verified with simulated data and through analysis of admixture among populations of diploid and tetraploid *Arabidopsis arenosa*.

## Introduction

Species are distributed across geographic ranges and potentially heterogeneous environments, and experience barriers to dispersal, leading to clinal genetic differentiation, genetic subdivisions into local populations or ‘demes’, or some combination of both (Endler, 1977; Bradburd *et al.*, 2013; Gompert & Buerkle, 2016). Even species with high rates of dispersal can have geographic ranges that are large relative to dispersal distances (e.g., Novembre *et al.*, 2008; Phifer-Rixey *et al.*, 2018), such that the distribution of traits and alleles is commonly heterogeneous and stratified among geographic locations. Quantifying this population heterogeneity and stratification is a fundamental component of empirical population genetics, both to provide a context for the study of evolutionary dynamics and as a component of learning about trait genetics in natural populations. Information about population structure and mixtures can reveal aspects of the underlying evolutionary processes and has played a significant role in shaping our understanding of the nature of hybridization, speciation, and adaptation. This includes knowledge of the prevalence of gene flow and introgression, as well as variability in introgression among geographic sites and genomic regions (e.g., Nadeau *et al.*, 2012; Abbott *et al.*, 2013; Gompert *et al.*, 2014b; Mandeville *et al.*, 2017; Meier *et al.*, 2017). For example, `structure`-like models are commonly used to quantify the proportion of an individual’s genome inherited from hypothetical source populations, which corresponds to their ancestry or admixture composition (Pritchard *et al.*, 2000; Falush *et al.*, 2003; Gompert *et al.*, 2014b). Beyond `structure`-like models, there is considerable interest in estimates of locus-specific ancestry and introgression, with a corresponding wealth of existing and continuously developing methods in computational statistics (e.g., Sankararaman *et al.*, 2008; Gompert & Buerkle, 2013; Gompert, 2016; Rosenzweig *et al.*, 2016; Ottenburghs *et al.*, 2016; Schumer *et al.*, 2019, for a review, see Gompert *et al.* 2017). These include parametric methods for detecting loci with ancestry that is concordant with the remainder of the genome (e.g., Szymura & Barton, 1986; Gompert & Buerkle, 2011a), or for detecting breakpoints and tracts of ancestry among chromosomal blocks or haplotypes (e.g., Wegmann *et al.*, 2011; Lawson *et al.*, 2012; Sohn *et al.*, 2012; Gompert, 2016). Similarly, researchers have contrasted ancestry and introgression of sex chromosomes relative to ancestry of autosomes in hybrid zones (Harrison & Larson, 2016; Chaturvedi *et al.*, 2020).

Accounting for population stratification and mixtures is typically a critical component of trait mapping in natural populations. Accounting for population stratification can reduce the number of false positive associations between loci and trait variation (e.g., Pritchard & Donnelly, 2001; Haworth *et al.*, 2019). Admixture coefficients or genetic kinship matrices can quantify diffuse genetic effects that are attributable to the genetic background of individuals (overall ancestry), rather than the effects of individual genetic loci (Zhou *et al.*, 2013; Hellwege *et al.*, 2017).

Despite the abundance of non-parametric statistical methods (e.g., `EIGENSTRAT`, Price *et al.* 2006 and `DAPC`, Jombart *et al.* 2010) and parametric models for population structure, methods for quantifying admixture in autopolyploid or mixed-ploidy individuals (combination of autosomes and sex chromosomes within individuals, or a mixture of ploidal levels among individuals in a population) are not fully developed. This is true even though 16% of all plant species contain some ploidal variation (Rice *et al.*, 2015). The dynamics of mixed-ploidy species can reveal processes governing polyploid evolution and the role of ploidal variation in adaptation and speciation (Kolář *et al.*, 2017).

Autopolyploids harbour multiple complete haploid subgenomes with sets of homologous chromosomes that share recent common ancestry and that aggregate and then segregate randomly in meiosis, leading to polysomic inheritance. Hence, methods for autopolyploid genetics should contain the ability to treat each allele copy at a locus as being independent.

In contrast, in allotetraploids with disomic inheritance, loci can be modeled as having diploid genotype values (and use the methods previously developed for diploids), instead of modeling complete tetraploid genotypes as with autotetraploids (even with minimal information on the origin of reads from the two different subgenomes using the model presented in Blischak *et al.*, 2017). `structure` can be used with autopolyploid and mixed-ploidy individuals, but lacks the ability to use genotype likelihoods as input data and thereby account for uncertainty in genotype calls, and requires a model misspecification to accommodate variable ploidy (i.e., by assuming a single ploidal level for input genotype data across all individuals, Meirmans *et al.*, 2018; Stift *et al.*, 2019). Differences in genotyping errors could occur across ploidal levels and cause potential artefacts if `structure` were applied to a mixed-ploidy data set of called genotypes, though the magnitude of such effects in estimation have not been well studied (Ferretti *et al.*, 2018). As a result, models that take genotype count data as input cannot make full use of low-depth sequencing, or be used as a population model for estimating genotypes (including imputation of missing genotypes), regardless of ploidal variation. However, other methods that use genotype likelihoods and low depth sequences have not been extended to polyploids (Skotte *et al.*, 2013; Meisner & Albrechtsen, 2018). The use of the full distribution of genotype likelihoods (from variant calling softwares like `GATK`, McKenna *et al.* 2010, `SAMtools`, Li 2011, or `FreeBayes`, Garrison & Marth 2012), rather than point estimates of genotypes, is particularly appropriate for polyploids in which a heterozygous genotype can arise from multiple dosages of alternative alleles (e.g., 1:3, 2:2, and 3:1 in a tetraploid) that will be difficult to distinguish, particularly with low sequencing depth. More generally, methods that use genotype likelihoods from all appropriately filtered loci will make more complete and better use of the available genomic data to estimate ancestry and genotypes (Gompert & Buerkle, 2011b; Nielsen *et al.*, 2012; Buerkle & Gompert, 2013; Vieira *et al.*, 2013), including for estimating genotypes to map phenotypes to the genomes of polyploids (Grandke *et al.*, 2016). In addition to the class of `structure`-like models for population allele frequencies and individual ancestry, methods have been developed to estimate genotypes from polyploid sequence data, without considering population structure and admixture (`EBG`, Blischak *et al.* 2017; `updog`, Gerard *et al.* 2018). These methods estimate genotypes by modeling typical sources of errors arising in high-throughput sequencing with additional information about the inheritance and segregation patterns in polyploids.

Similar to related approaches, the model presented herein (implemented in the software `entropy`) specifies genotype as a parameter that is informed by the genotype likelihood (the data), and by individual (admixture coefficient) and population parameters (allele frequency) that define the prior probability of observing a particular genotype. Our population model is suitable for updating genotype probabilities, which is particularly useful at genomic locations with low sequencing depth, or for computing genotypic probabilities at sites with no observed data (imputation). Point estimates or the full distributions of those genotype probabilities can then be used in downstream analyses. It has been shown that using an evolutionary model for allele frequencies in populations, including a `structure`-like model, improves estimates of genotypes from sequence data relative to methods that do not use population models (Gompert *et al.*, 2014b; Clark *et al.*, 2019).

A recent simulation study (Stift *et al.*, 2019) showed that model-based approaches like `structure` outperform other ancestry-estimation methods for the analysis of mixed-ploidy populations. At the same time, the assumption of `structure`-like models of admixture among ancestral demes should be tested, so as to avoid model misspecification and being misled for some populations and instances of gene flow (e.g., when there is additional sub-structure within the assumed ancestral populations, or inference is based on discrete samples along a continuous, isolation by distance gradient, etc., see Lawson *et al.*, 2018). Whereas the model can apply well to cases of contemporary hybridization and population mixtures, model misspecification can lead to incorrect inferences. Hence, the importance of model choice and fit has spurred further development of methods to gauge the appropriateness of the model for individual studies (Gompert *et al.*, 2014b; Garcia-Erill & Albrechtsen, 2019; Chaturvedi *et al.*, 2020).

With these motivations, we extend and thoroughly test the performance of a model similar to the admixture model implemented in `structure` (a version for diploids was presented previously as part of analyses in Gompert *et al.*, 2014b) to detect and quantify contemporary population structure in mixed-ploidy populations. In our `entropy` software, we specifically model mixed-ploidy by allowing for variable ploidal level across individuals (ranging from haploid to hexaploid). We have implemented methods for autopolyploids, since allopolyploids can be modelled as a lower ploidy, given sufficient knowledge of genome organization and chromosome pairing, including which loci occur on the pairs of homoeologous chromosomes (Bourke *et al.*, 2018). Herein we also restate a novel ancestry-estimation method for diploids that can detect and distinguish among early-generation hybrids using a parameter for the combination of ancestries at a locus (the *ancestry complement* model; previously presented in the Supplementary Material of Gompert *et al.*, 2014b). The *ancestry complement* model is an important extension to `structure`-like model for studies of early-generation hybrids, where, for instance, the presence of only first generation hybrids implies different dynamics than backcrosses and later generation hybrids (F2, F3, etc.). From our testing, we conclude that the `entropy` model has high accuracy rate in recovering true genotype and ancestry estimates from a variety of simulations, and further resolves population mixtures in empirical data from a diploid-autotetraploid hybrid zone.

## Methods

### Model specification

Our hierarchical Bayesian model describes the probability of parameters of interest (genotype, population allele frequency, admixture proportion, etc.) given the data (genotype likelihoods for individual SNPs), and is similar to the admixture model implemented in the software `structure` (Pritchard *et al.*, 2000; Falush *et al.*, 2003). Since the joint product across these hierarchical levels does not have a closed form, analytical solution, we rely on Markov chain Monte-Carlo (MCMC) methods to obtain samples from the posterior probability distributions of these parameters. Several related models have been implemented over the years that use a similar idea to obtain parameter estimates through various computational techniques, the most commonly used being Bayesian MCMC (e.g., Pritchard *et al.*, 2000) and Expectation-Maximization (EM) of a likelihood (e.g., Tang *et al.*, 2005), and more recently, variational inference of a posterior (e.g., Raj *et al.*, 2014). We chose to use Bayesian MCMC so as to obtain measures of uncertainty associated with the estimates of our parameters, especially since we wanted the model to be usable with uneven and low depth DNA sequence data. The measures of uncertainty are useful in interpretation of point estimates and can be carried forward into subsequent analyses.

We deviate from several previous models by using genotype likelihoods as input instead of fixed genotypes, as a way of propagating this uncertainty from the data to the inference of parameters. One can think of `entropy` as a data generative model that tries to match the genotype likelihoods (or genotypes) that are observed to an evolutionary process (parameterized by the allele frequency *p*, ancestry *z* and so on) that could have generated the data (Figure 1).

**Figure 1:**
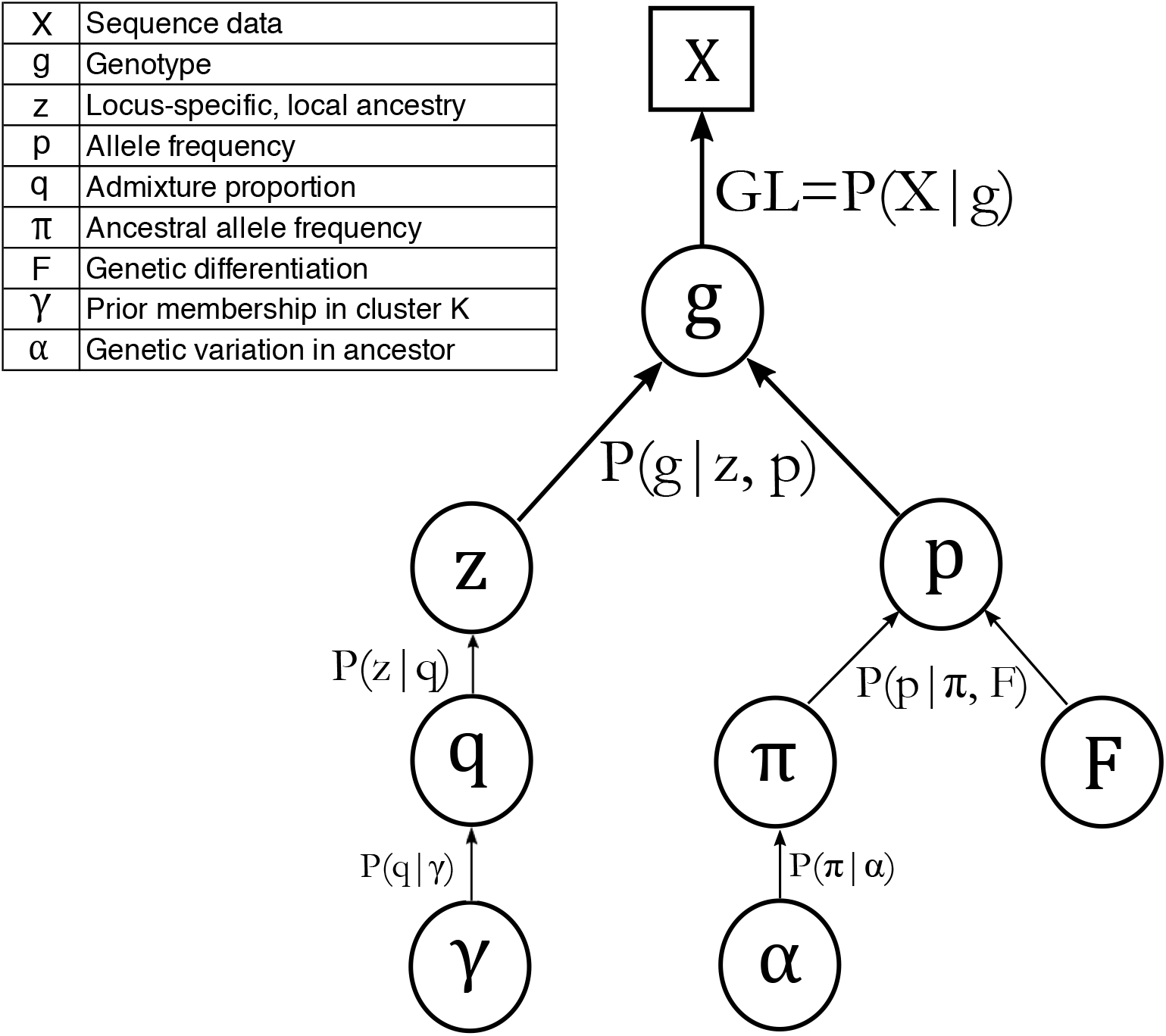
The graphical representation of the `entropy` model illustrates the information sharing in the model. Parameters that are being estimated are represented inside circles and the input sequence data are represented inside the square. The probability functions that generate these quantities are presented below each parameter. Typically, in a hierarchical framework we start at the bottom with new estimates of random values following a prior probability distribution. Then, conditional on these parameters, in the next highest level in the hierarchy we obtain estimates, and so forth, until the top level (the likelihood, here *GL* = *P* (*X* | *g*)) where estimates are constrained and estimated according to information in the data and prior probabilities. The parameter **q** is replaced by **Q** when using the *ancestry complement* model.

The evolutionary process we assume here starts with an ancestral population (characterized by allele frequency, *π*) that evolves through drift (parameterized by *F* using the Balding-Nichols model, Balding & Nichols, 1995) to give rise to the *K* ‘parental’ populations (each characterized by allele frequency, *p*) from which potentially admixed individuals are drawn, with admixture quantified by proportion *q* (Figure 1). We then use the observed genotype likelihoods (obtained from sequencing individuals) to match this evolutionary process and estimate our parameters through a hierarchical Bayesian model.

In the sections that follow, we describe each of the conditional probabilities involved in sampling and estimating these parameters of interest, moving from the base of the model hierarchy to the likelihood (Figure 1). A more detailed description of the mechanics of the MCMC procedure is provided in the Supplementary Material.

#### Admixture proportion (q and Q)

In this model, admixture proportion or ancestry in an individual is the proportion of an individual’s genome that is derived from one of *K* source populations. The admixture proportions are estimates of the average genome-wide or global ancestry for an individual and, with information on the individuals descended solely from parental populations, can be used to describe hybridization among the demes represented in the sample (as shown in Gompert *et al.*, 2017).

The `entropy` model includes two different models for the estimation of admixture proportion of individuals. The first is the *q*-model, similar to the `structure` admixture model with correlated allele frequencies (Falush *et al.*, 2003). Here, we specify a vector of admixture proportions of length *K*, **q**= [*q*_1_*, q*_2_*, …, q_k_*] to indicate the proportion of an individual’s genome that was inherited from each source population. These parameters are the probability of sampling a particular ancestry for an individual allele copy at the locus, independent of other alleles, which is equivalent to Hardy-Weinberg expectations that arise from random mating.

The second model in `entropy` is the novel *ancestry complement* model and considers the combination of ancestry for pairs of alleles across all loci in diploid individuals (the model is specified for diploids only). In early generation hybrids, interspecific (or interdemic) combinations of alleles are expected to be common. Parameterization of the ancestry combination of the pair of allele copies in this model allows for deviations from independence, which is assumed among allele copies in the simpler *q*-model. The ancestry combinations are represented in a *K × K* dimension matrix, **Q**. For example, with *K* = 2 demes the ancestry complement matrix **Q** is 2 *×* 2 in dimension. *Q*_11_ denotes the proportion of the individual’s genome in which both allele copies are descended from source population 1. Similarly, *Q*_22_ denotes the proportion of the individual’s genome in which both copies are descended from source population 2, and *Q*_12_ = *Q*_21_ denotes the proportion of the genome in which one allele copy is from source population 1 and the other allele copy is descended from source population 2 (since the order of the allele copy does not matter, *Q*_12_ is equal to *Q*_21_). In the *ancestry complement* model, the admixture proportion vector **q** is a derived quantity from the admixture complement matrix **Q**. For instance, with *K* = 2 demes, 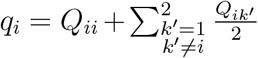 for *i* = {1, 2}.

As noted above, the benefit to the admixture complement parameterization is that it explicitly models the the combination of ancestry states at a locus, which is particularly beneficial in distinguishing among early generations of hybrid individuals (i.e., F1, F2, F3, and BC1). For first generation hybrids between parental taxa (F1) and between hybrids that have no parentage involving backcrossing (F2, F3, etc.) the expected value for *q*_1_ is 0.5, with some variance in observed individuals. This means that with the **q** vector alone, we can distinguish recent hybrids from the parentals and maybe backcrosses but not distinguish F1s from later generation hybrids. Likewise, distinguishing backcrosses from the parentals for later generations of hybrids is difficult with the admixture proportion vector **q** alone, given chance deviations from the expected values (Lindtke *et al.*, 2014). Particularly for early generations of hybridization between a pair of taxa, the combination of information in the admixture complement matrix **Q** (particularly *Q*_12_) and **q** can support assignment of individuals to hybrid generations (Figure 2).

**Figure 2:**
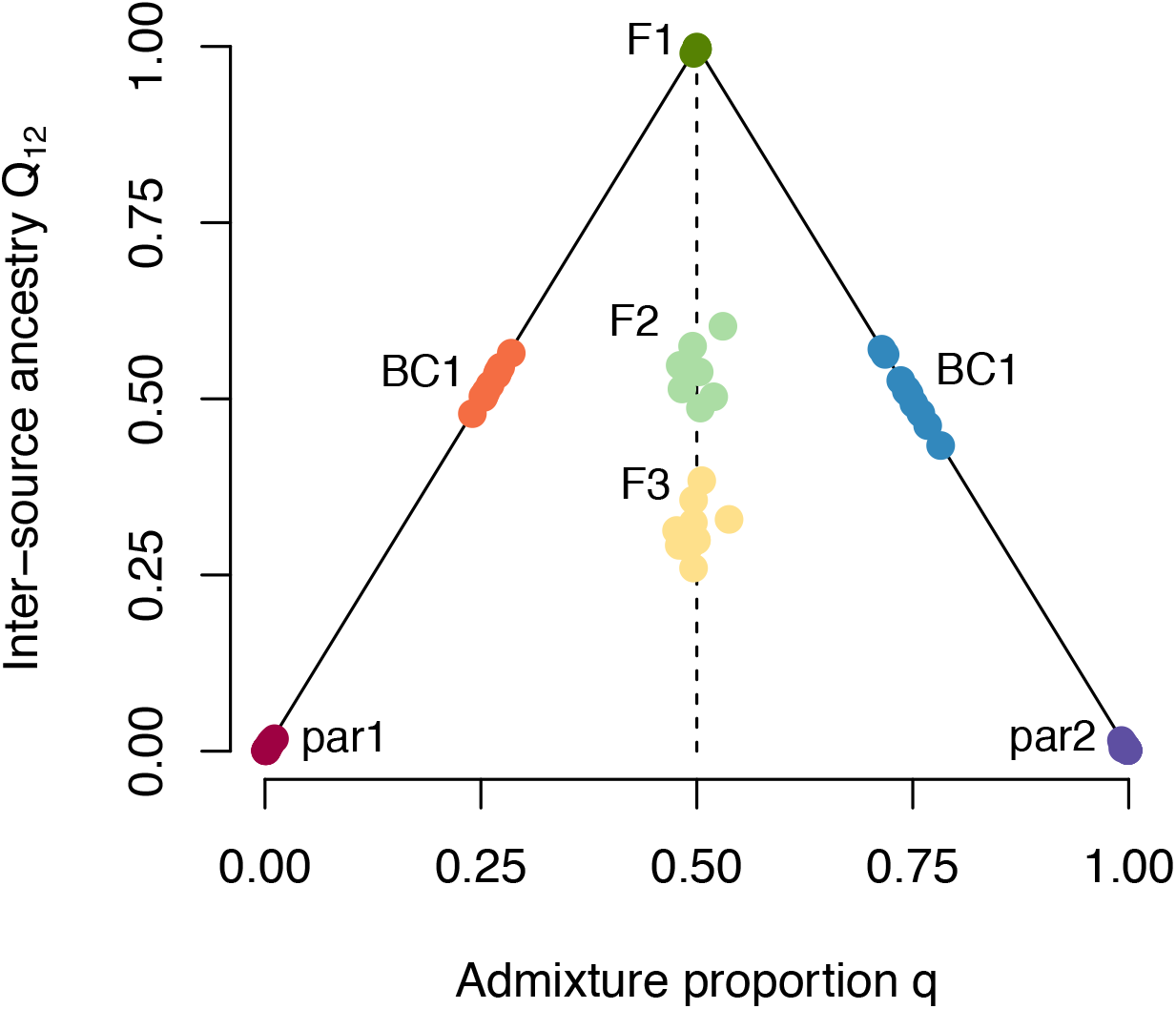
The diploid *ancestry complement* model in entropy with F1 hybrids (along the *q* = 0.5 line) between two parental populations (*K* = 2, at *q* = 0.0 and 1.0) reveals parameter differences between different early hybrid generations. Each color corresponds to individuals from a certain ancestry class: the parental species are colored in maroon and purple (at either ends), with the hybrids in varying shades from green to yellow and the back-crosses in orange and blue. The combination of admixture proportion *q* and ancestry complement **Q** distinguishes among F1, F2, and F3 hybrids. The admixture proportion (*q*) values for these three classes of ancestry are all 0.5 with some variance, but *Q*_12_ declines from 1 in the F1 with each generation of hybridization. Additionally, BC hybrids have maximal *Q*_12_ for a given *q*. The solid lines for the triangle indicate individuals with maximal possible *Q*_12_ values, corresponding to having at least one non-admixed parent.

Use of the admixture complement model will typically be restricted to low levels of *K*, because interpretation becomes increasingly complex for *K* > 2, requiring multi-dimensional plots for combinations of higher *K* values in the *Q* matrix (see Figure 2). In empirical study of systems for which *K* = 2 was well supported, the *ancestry complement* model has been used to learn about patterns of hybrid matings among *Lycaeides* butterflies (Gompert *et al.*, 2014b; Chaturvedi *et al.*, 2020), and *Catostomus* fish (Mandeville *et al.*, 2017, 2019), and in a related model used to study assortative mating among *Populus* species and their hybrids (Lindtke *et al.*, 2014). The *ancestry complement* model for diploids is not found in `structure`, or other `structure`-like models. As noted previously, the implementation of the *ancestry complement* model only was made in software for diploids, because the number of dimensions required to represent this matrix for higher ploidal levels was unwieldy and difficult to summarize into interpretable statistics.

#### Locus-specific, local ancestry (z)

The local ancestry parameter (*z*) is a marker for the population of origin of each allele copy at a given locus (for the **q** model; see below for the *ancestry complement* model) for an individual in a data set. It follows that the ancestry at a locus in an individual is informed by the genome-wide admixture proportions of that individual, reflecting different source populations, with *z* indicating the appropriate source population. So the prior probability for local ancestry of an individual *i* at locus *j* is given by the admixture proportion for that individual, *q_i_*: *P* (*z_ija_* = *k*) = *q_ik_* for allele copy *a* ∈ {1, 2, …, *n*} in autopolyploid individuals (since we model each allele copy to be independently derived). This allows for each allele copy at a locus to be derived from a different source population. This number is a single draw from a multinomial distribution conditioned on the admixture proportions in **q**.

The *z* vector for diploid individuals in the *ancestry complement* model functions similarly, in that we assume the conditional probability for local ancestry to be *P* (*z_ij_* = *kk’*|**Q**) = *Q_kk’_*. This means that the probability that both allele copies at a locus were inherited from source population *k* is equal to the proportion of the individual’s genome in which both allele copies are inherited from population *k*, and so on. This allows for the combinations of interspecific ancestry to be modeled explicitly as it considers the possibility of separate ancestry states at a locus.

#### Allele frequency (p)

The allele frequency in inferred demes is an important parameter that allows sharing of information among loci by quantifying their shared evolutionary divergence from allele frequencies in an idealized ancestral population (parameterized by *F* and *π*). The allele frequency in `entropy` for a locus *j* in population *k*, *p_jk_* is modeled with an F-model prior as in Nicholson *et al.* (2002); Falush *et al.* (2003); Gaggiotti & Foll (2010)

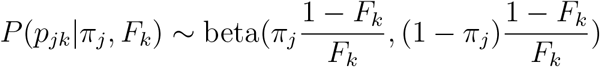

where *π_j_* denotes the allele frequency at locus *j* in the hypothetical population that was ancestral to the *K* source populations. *F_k_* denotes the extent to which the *k^th^* source population has diverged from the ancestral population. This is analogous to Wright’s *F_ST_* under all conditions of divergence and directly equivalent for an isolation model in which derived populations evolve due to drift relative to the ancestral population, with all loci in the genome sharing the same drift parameter (*F_k_*). This parameterization allows for a different *F_k_* value for each population relative to the inferred ancestral population (Nicholson *et al.*, 2002; Falush *et al.*, 2003). The prior on *π_j_* is beta(*α*, *α*) and the prior on *F_k_* is uniform(0,1), where *α* is inversely proportional to genetic variation in the ancestor and is estimated from the data. This formulation does not change for polyploid populations as is shown in the Implementation section of Nicholson *et al.* (2002).

#### Genotype (g)

In the `entropy` model the genotypes are treated as parameters and are estimated from the element-wise product of the genotype likelihood (the input data) and the prior probability for the genotypes, **GL**× *P* (*g_ij_*|*p_j_, z_ij_*). With contemporary DNA sequencers, genotypes are not observed directly, but instead information about these values is obtained through genotype likelihoods (**GL**) from bioinformatic steps and a model for the observed sequence data (incorporating read counts, base quality scores, mapping quality scores, etc.). Because these likelihoods are for discrete genotypes (count of alternate alleles at a locus), they can be readily rescaled so that they sum to one and can be used as a discrete probability distribution. Often, during the analysis of DNA sequencing data, software is used to call a genotype, for each locus and individual, to be the most likely genotype given the sequence data at the locus (i.e., the mode of the genotype likelihood). The use of genotype likelihoods rather than point estimates of genotype allows uncertainty stemming from sequencing depth and mapping quality to be incorporated into a probability distribution, while maximizing the use of information in sequence data. Genotype likelihoods can be obtained from most variant-calling softwares (e.g., all ploidal levels: `GATK` McKenna *et al.* 2010, all ploidal levels: `FreeBayes` Garrison & Marth 2012, or haploid-diploid: `SAMtools` Li 2011), which can take into account the base and mapping qualities, haplotypic information, along with read counts to estimate a likelihood for the genotype.

The prior probability of each genotype is calculated from the allele frequencies in the corresponding source population, as determined by the ancestry of the allele copy or the ancestry combination of a pair of alleles in the *ancestry complement* model. The prior probability for genotype allows imputation of genotype probabilities for sites that have no sequence data (i.e., missing data), or improvement of estimates from low-depth sites. Software for variant calling assigns sites with no data an equal likelihood for each genotype, but the population prior can be highly informative about the possible genotypes at a locus given the individual’s ancestry in the different source populations. The genotype prior probabilities for a *n*-ploid individual *i* at locus *j* is given as

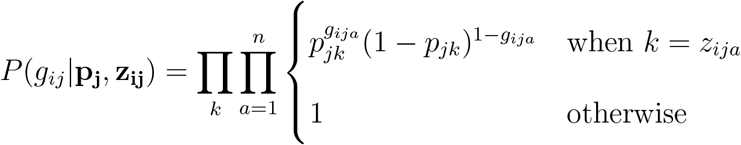

Here, **z_ij_** = [*k*_1_*, k*_2_*, …, k_n_*] denotes the local ancestry of the *n* allele copies for individual *i*, and *z_ija_* denotes the local ancestry of the specific allele copy, *a* in the individual. The term *p_jk_* denotes the corresponding allele frequency in the *k^th^* source population. The above expression yields a discrete posterior probability distribution of length *n* + 1 for each genotype (*g*) in a *n*-ploid individual ({0, 1, …, *n*}) i.e., number of possible alternate alleles at a locus, of the same size as the vector **GL**. Since we use a Bayesian framework for estimation, we obtain a discrete posterior probability distribution for genotypes, which can then be summarized by a point estimate to produce a continuous (mean or median) or integer (mode) value between 0 and *n*, the total number of allele copies at that locus. Point estimates can be the basis for further analysis with models that cannot use the full genotype probability distribution (e.g., input to other models and software).

### Model initialization and comparison

Given the potentially large number of loci and individuals in a contemporary study, the model will include large numbers of parameters, including *loci* × *individuals* genotypes (*g*), *loci* × *ploidy* × *individuals* × *populations* locus specific ancestries (*z*), *loci* × *populations* allele frequencies (*p*), and *individuals* × *populations* admixture proportions (*q*). Given the large number of parameters and Bayesian MCMC estimation, the efficiency of the estimation (faster convergence in this highly dimensional space) benefits from starting the chains as close to the stationary distributions as possible. Also due to the arbitrary nature of the model’s indexing of population or demes, estimation could include label switching among MCMC chains (i.e., the possibility of having ancestry or deme categories have different label indexes across chains, because the arbitrary indexing does not result in a change in the likelihood of the parameters given the data; see Stephens 2000). To speed convergence and avoid label switching, in practice one can initialize values based on a statistical procedure or taxonomic categories. We have used *K*-means clustering on the output of a linear discriminant analysis of the first five principal components (as specified in Jombart *et al.* 2010) to obtain estimates of the assignment probabilities to the *K* clusters for all the individuals. This analysis is run on point estimates of the genotypes from the genotype likelihoods. This statistical approach yields a probability of assignment of individuals to demes (the *K*-means clusters), without admixture. We have used the estimated assignment probabilities as mean initialization values (with some variance) for **q** in the `entropy` model and software (e.g., Gompert *et al.*, 2014b; Mandeville *et al.*, 2015; Haselhorst *et al.*, 2019). Additionally, starting values for the admixture proportions could come from taxonomic labels or justified strata in the sampling. The software implementation uses the initial *q* values to compute the initial population allele frequency in each of the *K* populations. This is calculated by finding the number of alleles with ancestry in a certain population (given by the initial **q**) and then dividing this number by the total number of allele copies in the population. This step initializes the population allele frequencies consistently among chains and limits the possibility of a label switch among chains.

Choice of an appropriate number of source populations, *K*, for a given set of data is a challenging problem, requiring consideration of both biology and statistics. Statistical methods for choosing the ‘best’ fit for a given model can be broadly classified into two categories: ones that provide measures for the likelihood of the observed genotype data given the parameters (for example, Pritchard *et al.* (2000); Alexander *et al.* (2009, etc.)), and ones that provide heuristic (*ad-hoc*) measures based on the ancestry estimates from the model (for example, Evanno *et al.* (2005); Raj *et al.* (2014, etc.)). The `entropy` model provides values of deviance (i.e., the negative log probability of the data given the parameters) and this can be used to calculate either the Deviance Information Criterion (Spiegelhalter *et al.*, 2002) or the Watanabe-Akaike Information Criterion (Watanabe, 2010). Consequently, a lower value for the information criterion signifies a better fit.

The summary of model fit, in combination with graphical analyses of **q** estimates, can contribute to an understanding of the number of potential demes (*K*) involved in admixture, particularly for taxa with contemporary hybridization and in the context of other information about the evolutionary history of the groups. However, this measure of model fit only allows contrasts among models for different choices of the number of demes (*K*). As such, `structure`-like models cannot themselves provide evidence for demic population structure (as noted in Pritchard *et al.* 2000), but instead must rely on complementary analyses and knowledge of the system. If the true population histories differ significantly from the underlying demic model, contrasts of the information criteria for different *K* can indicate which model best approximates the system. However, all of the demic models could fit poorly if genetic differences among individuals include substantial isolation by distance, additional substructure within the ancestral populations, rather than, or in addition to differences in the actual number of demes (*K*). Additionally, inference of the number of demes using `structure`-like models (or ordinations) can be misled by uneven sampling of individuals from putative demes, since very uneven sampling can introduce spurious substructure and lead to an underestimation of ‘true’ number of subpopulations (Puechmaille, 2016). Finally, aside from the difficulty of inferring the number of demes that are consistent with the data, the choice of *K* does not affect the estimation of genotypes. Instead, genotype estimates can be averaged over the posterior distributions of genotypes across all *K* runs to obtain a point estimate at a given locus for an individual (e.g., Gompert *et al.*, 2014b).

### Model performance

Our measures of model performance build on previous testing of the model and software for diploids (Gompert *et al.*, 2014b) and emphasize tests of the model extensions to simulated and empirical data from polyploid and mixed-ploidy samples. As noted above, the diploid portion of the model has been used previously for several empirical analyses (e.g., Gompert *et al.*, 2014b; Mandeville *et al.*, 2015; Chaturvedi *et al.*, 2020). We test the performance of the *ancestry complement* model for diploid populations under similar simulation conditions. We point the reader to previous empirical studies (first in Gompert *et al.* 2014b, and subsequently in Mandeville *et al.* 2017; Chaturvedi *et al.* 2020, etc.) for further applications of the model to better distinguish among different classes of early generation hybrids and to distinguish recent from more advanced generation hybrids in diploid individuals (as seen in Figure 2).

#### Simulated data

We used simulations to quantify the performance of the model using three different metrics: accuracy in genotype and ancestry estimates under various simulation parameters (for each 2n, 3n, 4n, 6n, and 2n-4n data set), ability to impute missing data under varying missingness percentages (for each 2n, 3n, 4n, and 6n data set), and accuracy in ancestry estimates for a trade-off between coverage and sequence depth (4n data set).

The genotypic data for 2,000 loci and 100 individuals were simulated using the following evolutionary history. Individuals were assumed to be descended (either completely or partially) from one of *K* = 3 demes. The demes were a result of evolution with drift relative to an ancestral population, with the ancestral allele frequency at each locus drawn from a beta(0.5, 0.75) distribution to simulate the allele frequency spectrum expected in a real population with an abundance of low-frequency alleles. Separate simulations considered different amounts of evolution relative to the ancestral population, using an F-model for derived allele frequencies and *F* ∈ {0.05, 0.1, 0.2, 0.4} (ranging from low to high evolutionary divergence), and the differentiation this induced among demes. Based on the allele frequencies in demes, genotypes were simulated from a binomial distribution given the individual’s ploidy and their local ancestry across different populations. For instance, in a tetraploid individual, the genotype at a locus was drawn from binomial distribution with four draws (number of allele copies) and the success probability being the frequency of the alternate allele, weighted by its proportional ancestry in the source population. An individual could either be a parental, F1, back-cross between an F1 and a parental (BC1), F2, or F3. The genotypes were then converted to genotype likelihoods based on a range of sequence depths (drawn from a Poisson distribution with means: 1×, 2×, 4×, 6×, and 12×) following the GATK HaplotypeCaller model (Poplin *et al.*, 2017), assuming a constant sequencing and mapping quality so as to isolate any bias in these numbers on estimating our parameters. Typical sources of variance in sequencing were captured by simulating a Poisson random variable for depth at a site, including sites with very low depth or no sequence reads. Low depths correspond to greater genotype uncertainty, as reflected in the simulated genotype likelihoods.

Similarly, to explicitly validate the mixed-ploidy portion of our model, we simulated genotypic data for a hundred individuals, with fifty tetraploids and fifty diploids. Here, the genotypic data for the loci were drawn from the same evolutionary process as stated above with a change in the binomial sampling to yield the correct number of allele copies. The simulations were run for *F* ∈ {0.05, 0.1} and for an average sequence depth of 2× for diploid individuals and 4× for tetraploid individuals. As input, we also provided the ploidy of each individual along with the genotype likelihoods to `entropy`. The goal here was to primarily test the ability of our software to handle mixed-ploidy input and secondarily, to test the minimal ability of our model to recover the simulated parameters. Therefore, we chose a reasonable value of sequencing depth for tetraploids (i.e., 4×) since we knew the behavior of the model under lower depths from other simulations in this study. From these simulations, model performance was quantified by calculating the accuracy in estimation of genotype and ancestry across all individuals and loci.

One measure of model performance would be the extent to which the model could correctly impute missing (i.e., left-out) data from a simulation. To quantify the ability of the model to impute left-out data, subsets of the genotypic data from above were randomly excluded from a complete data set to achieve varying proportions of missingness (10%, 20%, 30%, 40%) over loci and individuals. This metric was important to test the performance of the model, not only for assessing the accuracy in imputing missing values, but also to mimic real empirical sequencing in which regions of the genome are not sampled at all for a number of individuals. This form of testing is akin to conducting a posterior predictive check in a Bayesian modeling framework (first introduced in Rubin, 1984) by quantifying the ability of our model to recover simulated parameters, especially from held-out (in our case, missing) data. Secondarily, this test of performance gives an indication of how data missingness would affect inferences of genotype and admixture proportions with empirical data, and we considered a range of missingness that one might encounter in empirical studies. To simulate this missing data, we randomly selected loci to have equally probable genotype likelihoods (i.e., every dosage/genotype is equally likely given no other information) to mimic the absence of sequence information at this locus.

To test the hypothesis that it is better to estimate average genome-wide ancestry by capturing more of the genome (i.e., loci, via greater genome coverage, defined here to mean extent of the genome covered by the sequence data) at a lower sequencing depth than it is to sequence a smaller region of the genome (i.e., lower coverage) at a higher depth (i.e., more reads), we ran a simulation for 100 tetraploid individuals across three pairs of values for coverage and corresponding sequence depth. The assumed evolutionary process was the same as the one used before, in which we simulated the tetraploid individuals as being descended from one of 3 possible demes (differentiated by *F* = 0.05) sequenced at: 4× and 1,000 loci (‘low’ coverage), 2× and 2,000 loci (‘medium’ coverage), and 1× and 4,000 loci (‘high’ coverage). Our testing here focuses on the “middle” tetraploid case, since we expect a similar mechanism to be operating at lower and higher ploidal levels (Buerkle & Gompert, 2013).

In total, 1,210 simulations were run to quantify accuracy in estimation, ability to impute missing data, and a trade-off between coverage and sequence depth across different levels of ploidy (2n, 3n, 4n, 6n, and 2n-4n), range of missingness percentages, varying levels of sequence depth, and admixture from three ancestral populations (at varying levels of evolutionary divergence). For simulations that contained missing data, we used the correlation metric between the point estimate of our parameter (the average of posterior distributions across chains) and the simulated truth to measure how well we could recapture this simulated parameter given a certain percentage of missing data. For the rest of the simulations, we calculated the root mean squared error (RMSE) between the inferred values for the genotype and admixture proportions and the known true values that were simulated, as a way of measuring our ability to recover the truth.

#### Empirical data

As a further test of the performance of the model and software, we reanalyzed an empirical mixed-ploidy data set that includes DNA sequences of individuals from diploid and autotetraploid populations of *Arabidopsis arenosa* across Europe (Monnahan *et al.*, 2019). We compared estimates of admixture proportions from this mixed-ploidy sample from both entropy and structure softwares. We used, as input, sequence data obtained from the vcf files for eight scaffolds, which were shared by the authors of Monnahan *et al.* (2019). From these, we sampled single variable loci randomly in 50,000 base pair windows (within each scaffold) to retain loci that were more likely to vary independently due to recombination and independent evolution. This left us with a set of 5,655 loci across 287 individuals (105 diploids in 15 populations and 182 tetraploids in 24 populations) with 22.4% missing data. The previous analysis in Monnahan *et al.* (2019) removed sites with excess heterozygosity and selected for variants that passed through the GATK Best Practices workflow (Van der Auwera *et al.*, 2013), to end up with 9,543 loci across 287 individuals with 2.4% missing data. Because we thinned loci randomly in 50,000 base pair intervals, without regard for missingness, our subset of loci reflected the average missingness among variants in the original vcf file, rather than the subset that was selected for low missingness (among other criteria) in Monnahan *et al.* (2019). We note that this process of random thinning only affects the credible intervals, and not the point estimates of the admixture proportions from the model. By including more loci in our analysis, we would get an incrementally more accurate point estimate for our genome-wide admixture proportion with a tighter credible interval. However, we also note that there is a diminishing return to including more loci in the analysis if the goal is to simply obtain point estimates for admixture proportion. The input to structure (version 2.3.4) was a file with the called values of genotypes (GT field in the vcf file) of selected loci and individuals, called using `GATK HaplotypeCaller` (version 3.5, McKenna *et al.*, 2010). Given that the maximum ploidy included four allele copies, the loci for the diploid individuals were encoded with four allele copies and the extra two allele copies as missing data (since the `structure` manual indicates that all individuals in the sample should have a single ploidal level, Meirmans *et al.*, 2018). The input to `entropy` was a file with the genotype likelihoods (PL field in the vcf file) of the selected loci, rather that the point estimates of the genotypes. The `entropy` model was initialized using the discriminant function method described previously, to reduce the chance of label switching among chains and speed MCMC convergence. We compared admixture proportion estimates from `structure` and `entropy` primarily for *K* = 6, which was regarded by Monnahan *et al.* (2019) as the most likely model given other knowledge of the evolutionary history of *A. arenosa*.

The `structure` admixture model was run three times for 600,000 iterations in total (which included a 100,000 iterations for burn-in), which took approximately 102 hours each. This number of iterations was chosen based on multiple runs of different lengths and picking the shortest run that arrived at approximately the same estimates as the longer runs. Since `structure` stores every sample after burn-in as a draw from the posterior distribution, the admixture proportions were estimated based on 500,000 draws. On the other hand, the `entropy` model was run with three chains simultaneously for 30,000 total iterations with 10,000 burn-in each, which took approximately 24 hours in total. The number of steps was chosen based on the convergence of previous data sets of similar size. The quicker convergence times were likely a result of starting our chains with plausible initial admixture proportions (as mentioned in the Model initialization and comparison section). Researchers could typically use `fastStructure` (Raj *et al.*, 2014) in this case, but this software only allows for diploid samples, which requires a downsampling at each tetraploid locus to fit the requirements for the input data (Monnahan *et al.*, 2019). The samples collected were thinned to retain every 10^*th*^ step to remove autocorrelation within the chain. We were finally left with 6,000 (2,000 ×3) samples from the posterior distribution for admixture proportion. The chains were tested for convergence by looking at the trace plots (to check for sufficient exploration of parameter space) and the average 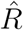 statistic (*≈* 1.01) across parameters (Gelman & Rubin, 1992). To validate the number of clusters statistically, we ran the entire *A. arenosa* data set for values of *K* ranging from 4 through 10 to obtain WAIC estimates, and compared them to estimates from Monnahan *et al.* (2019).

## Results

### Simulated data

We present the effect of sequence depth, *F*, number of ancestral demes, and ploidy on the ability of our model to accurately predict parameter estimates from simulated data (Figures 3 and 4). Based on the different axes of variation in our simulation parameters, we found that sequence depth had the strongest effect on our ability to estimate both genotypes and admixture proportions (for both models presented above) accurately, followed by the degree of differentiation *F* between our simulated demes. From our simulations containing missing data, as expected, the model performed better at recapturing the missing genotypes when we had lower percentages of missing data (Figures S1 and S2). Holding all else constant, we also found that we did better at accurately estimating admixture proportions of tetraploid individuals when we had higher coverage (number of loci across the genome) over higher sequencing depth (read depth at a locus; Figure 5).

**Figure 3:**
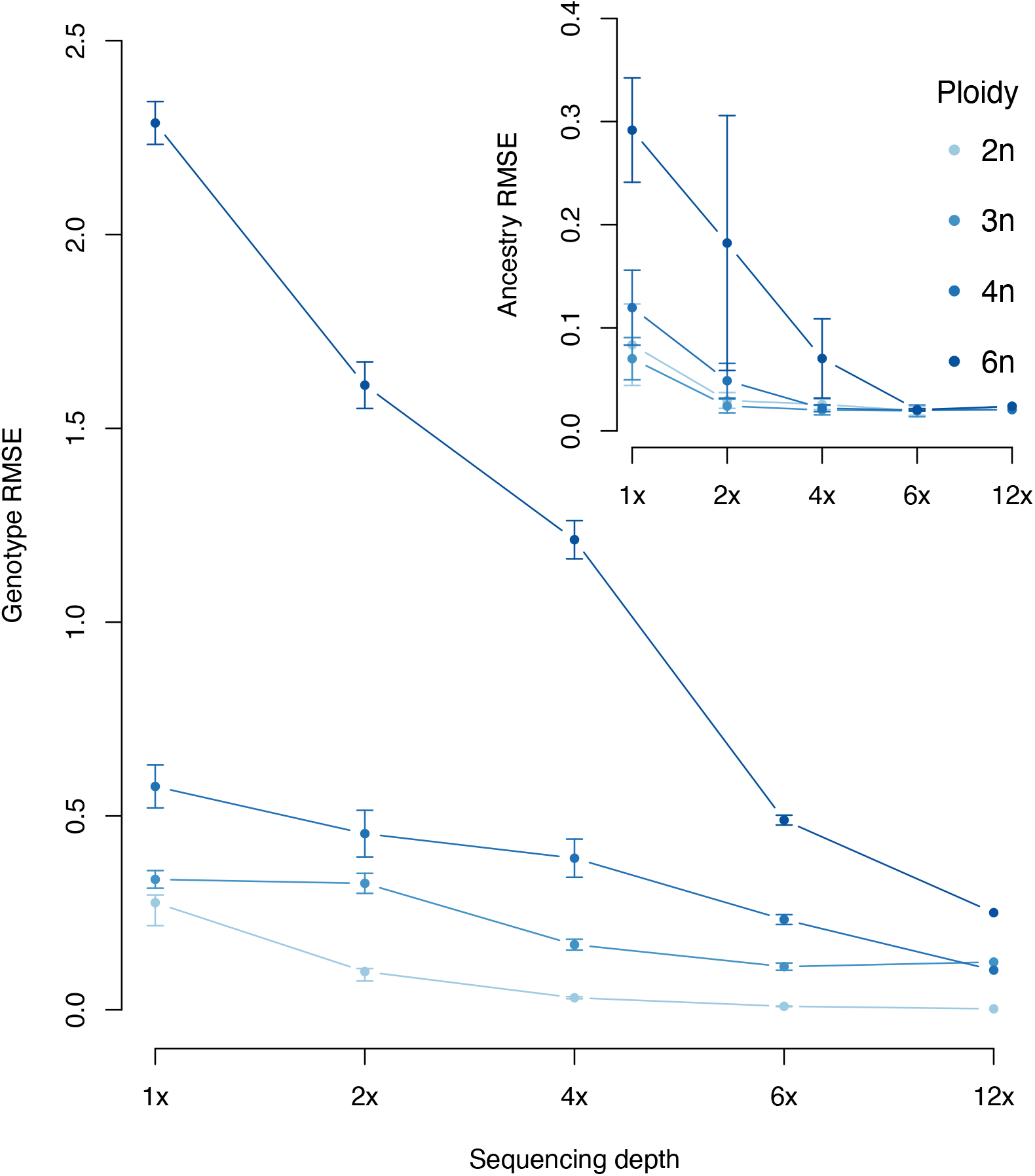
Error in genotype and ancestry estimation decreases with increase in sequencing depth across a range of ploidy and simulation parameters. The outer plot depicts declining RMSE in estimates of genotype for different ploidal levels with increasing sequence depth, across *F* = {0.05, 0.1, 0.2, 0.4} and number of source populations *K* = {1, 2, 3}. We found consistently higher error for lower sequence depth (across all ploidal levels). The inner plot depicts a similar decline in RMSE for estimates of admixture proportion for different sequencing depth and ploidy (with the same simulation parameters as before). The larger error bars at sequencing depth of 1× and 2× for higher ploidal levels is expected, but we see a levelling off at sequencing depths greater than 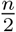×, where *n* is ploidal level, indicating high accuracy in admixture proportion estimation.

**Figure 4:**
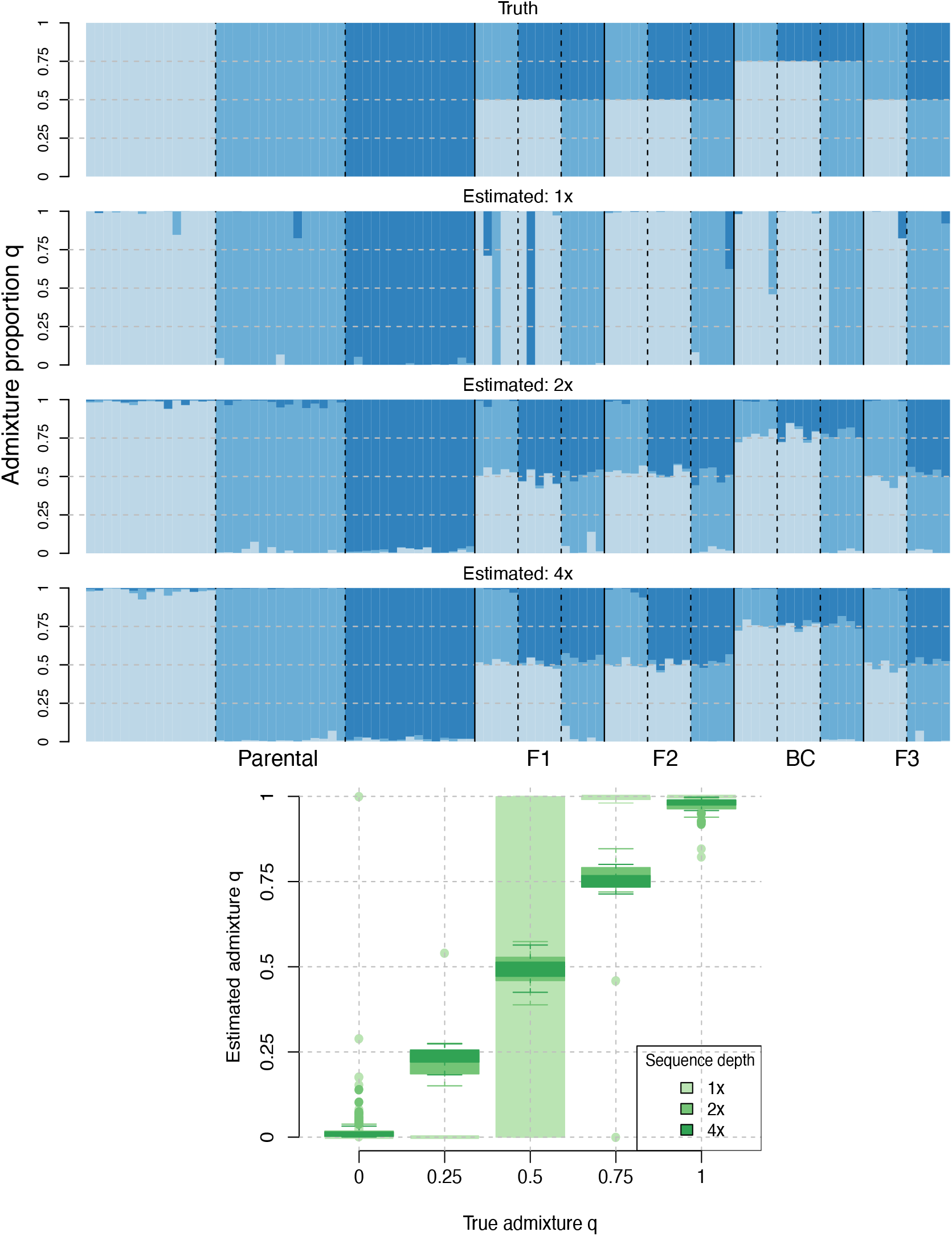
With increasing sequencing depth (rows 2–4, blue) the model more accurately estimates true admixture proportions (row 1, blue), particularly among hybrids (shown here for **tetraploid** individuals). With 1 × average sequence depth, `entropy` accurately estimated ancestry of parentals but did much less well with hybrids (≈(0, 1) for *q* = (0.25, 0.75)). With an average of 2× sequence depth at a locus `entropy` more accurately estimated admixture proportion, and at 4× average sequence depth estimates are very close to the truth. The second plot (green) illustrates the diminishing returns with higher depth, averaged across all ancestry groups. This plot contains comparisons of estimated and true admixture proportions for varying sequence depths across *K* = 3 source populations and 100 individuals and 2000 loci.

**Figure 5:**
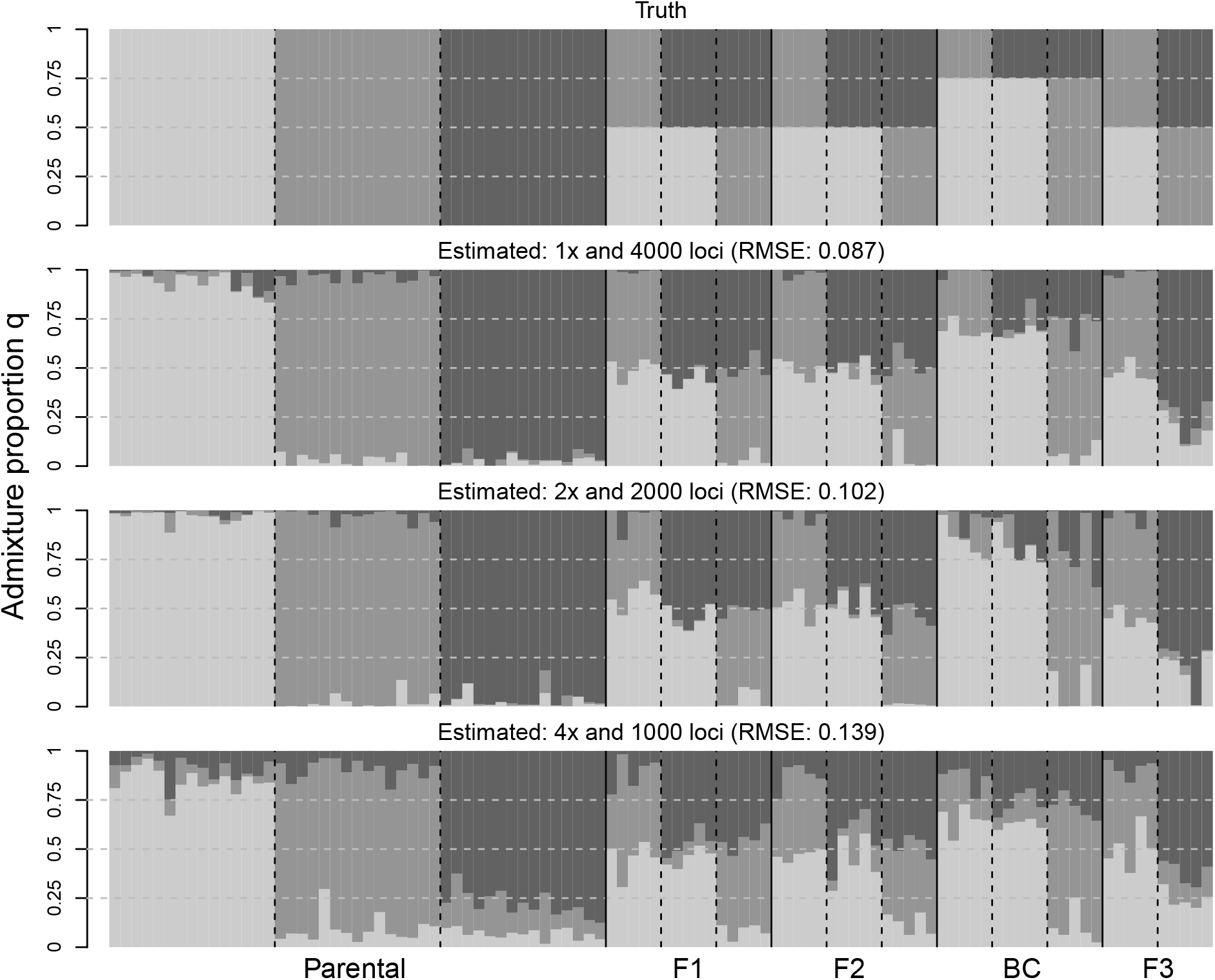
Admixture proportion is more accurately estimated with higher coverage and lower sequence depth. With higher coverage (i.e., more loci across the genome, 4000 loci) and a lower average sequence depth (1×), the estimates are closer to the simulated truth for global ancestry over the same data set subsampled to lower coverage (1000 loci) and a correspondingly higher sequence depth (4×). Admixture proportion estimates and RMSE values shown here for a continuum between 1× and 4× average sequence depth, with a corresponding coverage between 1000 loci and 4000 loci.

With regard to the estimation of genotypes, we better distinguished discrete genotype classes at higher sequence depth and higher *F* values. The reason we obtained larger values of RMSE for higher ploidy is a consequence of a wider range of genotypes that are possible. So as a consequence of the RMSE statistic, we are bound to get higher error values for higher number of genotype classes (i.e., higher ploidy) in our data. But, based on the approximately constant correlation between simulated and estimated genotypes in our missing data sets, we show that higher ploidal levels do not translate to a higher error rate in estimation (Figures S1 and S2). However, the spread around the RMSE across each ploidal level suggests that the degree of differentiation (*F*) and number of ancestral demes did not play a major role in our accuracy of prediction, as shown by the inset plot in Figure 3 and Figure S7.

Similarly, in the estimation of admixture proportions we found that the sequence depth had the biggest effect on our ability to recover the truth, followed by the degree of differentiation between the simulated demes, for both ancestry models presented here (Figures 4(a), S4, S5 and S6). With a sequence depth of 1×, we only observed one allele in a tetraploid and, as a result, our genome-wide ancestry estimates were solely guided by a single allele at each locus. However, the model performed much better once we observed, on average, two alleles (2×) at a given site and hit diminishing returns in sequencing beyond 4× (as seen in the estimates from Figure 4).

For the simulations involving missing genotype likelihood data, we found that the correlation (*r*) between the estimated genotypes at the missing sites and the simulated truth was between 0.76 and 0.88 across ploidal levels, indicating an increasing correlation with a decrease in missing percentage (Figure S8). This correlation translates to the fact that, on average across all simulation parameters, we could predict approximately 70% of our missing genotypes accurately (coefficient of determination, expressed as a percentage, is equal to *r*^2^). We also found that we have a higher correlation of estimated and true genotypes for higher ploidy levels, given the same missingness percentage and degree of differentiation (Figure S1).

Similarly, for admixture proportion estimation, we found a correlation between 0.96 and 1 for 296 out of 320 simulations. The set of outlier simulations were for low *F* = 0.05 and a high missingness (40%) value across all ploidal levels, for which the correlation was only 0.83 (Figure S3). For more than 80 percent of simulations, we predicted approximately 95% of our missing admixture proportions accurately (Figure S9).

Based on the RMSE metric, we found that the model estimated genome-wide ancestry for tetraploid individuals more accurately with higher coverage across the genome and a lower sequencing depth (4,000 loci at 1×) than with lower coverage and a higher sequencing depth (1,000 loci at 4×) (Figure 5).

### Empirical data

Overall, the admixture estimates from `entropy` closely match the estimates from `structure` (Figure 6). Similarly, the `entropy` and `structure` admixture estimates largely match those presented in Monnahan *et al.* (2019), which used a combination of analyses from `fastStructure` (Raj *et al.*, 2014) and a non-parametric K-means clustering technique (with a confirmatory analysis in `structure` for *K* = 6). Below we note some of the differences that were found between the ancestry estimates from `entropy` and `structure` (and shown in Figure 6).

**Figure 6:**
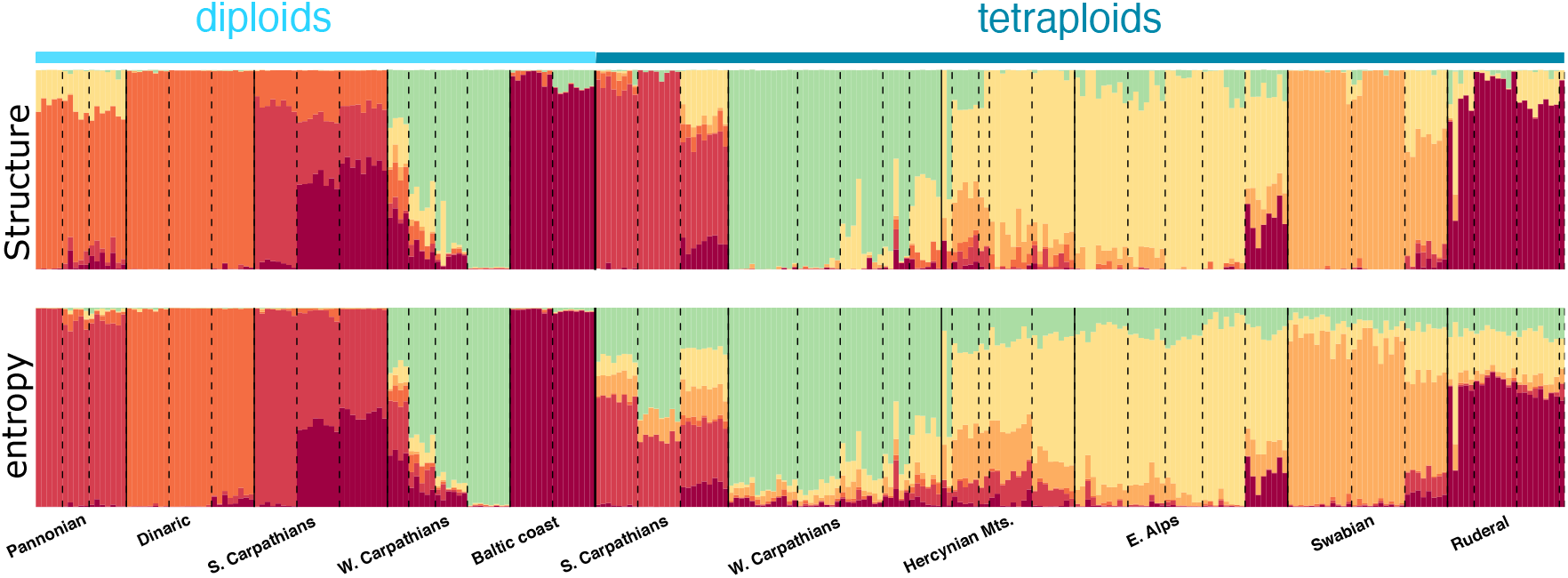
Admixture proportion estimates from structure and entropy agree very well for most the 287 *Arabidopsis arenosa* individuals from a mixed diploid and autotetraploid sample of populations across Europe for a *K* = 6 model with a median sequence depth of 10× (data from Monnahan *et al.*, 2019). The *K* = 6 model was the preferred model in the analysis by Monnahan *et al.* (2019). The two most notable differences between the `structure` and `entropy` estimates were the labeling of the Pannonian individuals (far left) as red ancestry by `entropy` versus a mixture of orange & yellow by `structure` and the different contributions to the composition of diploid and tetraploid S. Carpathians in the `entropy` and `structure` analyses.

Firstly, the admixture proportion estimates for the diploid individuals in populations from the Pannonian region were calculated to be different by `entropy` and `structure`. The `entropy` model assigned these individuals to a separate cluster, but the `structure` model found these same individuals to be genetically intermediate between the Dinaric (orange) and E. Alps (yellow) regional ancestries. Based on the evolutionary history of the plant, the Pannonian populations are the most divergent and should separate out as their own cluster (as shown in Figure 1 of Monnahan *et al.*, 2019). This hypothesis was further supported when both `entropy` and `structure` placed the Pannonian population into a distinct cluster when run for a *K* = 5 model on only the diploid individuals and a *K* = 7 model on all the *A. arenosa* individuals (as shown in Figures S10 and S12 for `entropy` and Figure S5 of Monnahan *et al.* 2019 for `structure`), as expected when running the analysis with a higher *K* in `structure`-like models. Secondly, the tetraploid individuals in the S. Carpathians are estimated to share some ancestry (between ~ 20% and 50%) with their W. Carpathian (green) counterparts in `entropy` but this was not found to be the case with the estimates from `structure`. This shared, intermediate or hybrid ancestry in the S. Carpathian populations is to be expected from the single origin of tetraploidy in the populations of the Carpathian mountain range, as confirmed through coalescent simulations of this mixed-ploidy hybrid zone (as presented in Figure 4 and Figure S9 of Monnahan *et al.*, 2019).

From our range of runs for *K* = 4 to 10, we found the lowest WAIC value for *K* = 9 (Table S1), as opposed to *K* = 6 that was found by Monnahan *et al.* (2019). The authors of the original study used a combination of the Bayesian Information Criterion (BIC, Schwarz *et al.* 1978), and the similarity index proposed by Nordborg *et al.* (2005) to inform their choice of *K*. However, the range of BIC values for *K* = 5 to 10 were found to be within five points of each other, indicating similar support for the given cases of *K*, highlighting the potential challenge of choosing the ‘right’ *K* in empirical studies.

## Discussion

In the context of recent hybridization or admixture among divergent lineages, estimates of admixture proportions are a fundamental component of analyses of evolutionary processes or of learning the effects of population stratification in genome-wide association studies (Gompert & Buerkle, 2013; Harrison & Larson, 2016; Gompert *et al.*, 2017). Hybrids commonly occur between taxa that have sex chromosomes (and mixed-ploidy within genomes of the heterogametic sex) and sex chromosomes may contribute disproportionately to their reproductive isolation (Payseur *et al.*, 2004; Sæther *et al.*, 2007; Presgraves, 2008; Macholán *et al.*, 2011; Chaturvedi *et al.*, 2020). Additionally, many species complexes involve interactions and potential hybridization between individuals of different ploidy, including autopolyploids (e.g., Otto & Whitton, 2000; Kolář *et al.*, 2017; Van de Peer *et al.*, 2017). Population genetic analyses of polyploids would benefit from models that correctly specify the number of allele copies at a locus, rather than misspecified models that do not fully use the available data (e.g., encoding diploids as tetraploids with missing data so that structure can be used to analyze mixed ploidy individuals). Additionally, given genotype uncertainty in contemporary low-depth sequencing data from populations, we make better use of the data with models that formally incorporate uncertainty through the use of genotype likelihoods as input (Buerkle & Gompert, 2013; Fumagalli *et al.*, 2013). Here, we address these needs and present additional benefits, with improvements in running time, and ability to assess convergence of chains using appropriate metrics, in the form of a population model for allele frequencies and admixture of individuals that follows the precedent of the `structure` model (Pritchard *et al.*, 2000; Falush *et al.*, 2003) and its several derivatives. We present and analyze the performance of `entropy`, a hierarchical Bayesian model that can use genotype likelihoods to estimate genotype and ancestry for polyploid and mixed-ploidy individuals.

We found that the `entropy` model performed well to capture the truth from simulated mixed-ploidy data sets. We used estimates from the model for simulated autopolyploid data and quantified similarity of our estimates to the known values using RMSE and correlation statistics. The `entropy` software implements a population model and information sharing among individuals and loci that provides stronger evidence for low-depth, or missing genotypes (especially with polyploids and low-depth sequencing) than methods that do not model populations (consistent with Clark *et al.*, 2019). With the extension of the model and software to mixed-ploidy, we can also model haploid loci or hemizygous regions of the genome. Thus, given knowledge of the genomic position of loci, the model will support contrasts of ancestry between sex chromosomes and autosomes (e.g., Hamilton *et al.*, 2013; Parchman *et al.*, 2013; Harrison & Larson, 2016). For the analysis of diploid hybrids, as shown previously in Gompert *et al.* (2014b), the *ancestry complement* model considers the combination of ancestry in diploid genotypes and allows genotypic data to more readily distinguish among different classes of early generation hybrids (Figure 2) and to distinguish recent from more advanced generation hybrids (e.g., Gompert *et al.*, 2014b; Mandeville *et al.*, 2017; Chaturvedi *et al.*, 2020).

In our simulations, we found that sequence depth had the largest effect on accurately estimating the genotype and admixture proportion of an individual for a given ploidal level, similar to findings in Gerard *et al.* (2018). The degree of differentiation among demes (driven by *F* divergence from the ancestral population) had the second largest effect on the accuracy of genotype and ancestry estimates (as seen in Figures S2, S4 and S6, and Table S2). For admixture proportion *q*, we found no difference in our ability to estimate admixture proportion across the different ancestry classes (F1, F2, BC, etc.; as seen in Figure S4). Across our simulations, we also found that the percent of missingness did not affect how well we could estimate true parameters. For example, when going from 40% to 10% missingness in sequence data there was only a 2% gain in accuracy of prediction for tetraploid genotypes. In summary, from the different combinations of the simulation parameters, it was the hardest to recover parameters accurately when we had low sequence depth (for higher ploidy) and minimal differentiation between populations (*F <* 0.05), as was expected. Based on the ability to recover the truth in various simulations, for analyses of admixture proportions with this model we recommend choosing a median sequence depth of 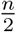× (i.e., 2× for tetraploids) and sampling more individuals and populations rather than sequencing deeply (consistent with findings from our simulations in Figures 4(a) and (c) and Buerkle & Gompert, 2013; Fumagalli *et al.*, 2013). However, if the goal of an analysis is highly accurate genotype estimates, as expected, sequencing to 6× or greater depth might be warranted (Figures 3 and S5).

Our direct comparison of `entropy` estimates to estimates from `structure` for an empirical mixed-ploidy data set (diploid and tetraploid) of *Arabidopsis arenosa* (Monnahan *et al.*, 2019) validated the software implementation of the model and revealed some differences of admixture proportions and inferred ancestry for a few populations. The data used with `entropy` and `structure` contained fewer loci (5655 loci versus 9543 loci) and a higher percentage of missing data that is typical in a RADseq or similar dataset (22.4% versus 2.4%), compared to the original analysis in Monnahan *et al.* (2019). Nevertheless, the two models were still able to capture the previously inferred population structure. Even though the median sequencing depth of individuals across this study was 10×, we have no reason to believe that ancestry estimation would be affected by lower sequencing depth. Based on a posthoc analysis of the posterior probabilities we found no clear association between individuals with low sequencing depth and higher uncertainty in ancestry estimates. This trend was true across populations and within individuals. For example, the WEK population had the lowest median sequencing depth at 6× (with certain individuals in the population averaging 2×), however it was placed in the third quartile when ranked on decreasing standard deviations for the admixture proportion estimates. The estimates from the `entropy` model for a cluster of admixed tetraploid individuals indicated a portion of their ancestry belonged to a previously undetected neighboring cluster, a finding that was supported by coalescent simulations in Monnahan *et al.* (2019), but was not captured by the `structure` model. Additionally, the `entropy` model distinguished and assigned some exceptional individuals to a distinct cluster instead of classifying them as belonging to an admixed group, as done by `structure`. The support for a higher number of source populations (WAIC supported *K* = 9, in Figure S13, as the ‘best’ fit for the data, rather than the *K* = 6 from the original manuscript) stemmed from a distinct clustering of diverged individuals and a finer subdivision in ancestry of individuals in the mixed-ploidy hybrid zone. However, as noted previously, the interpretation of population subdivision into *K* clusters requires support from additional evidence (see Tables S3 and S4), as was done in Monnahan *et al.* (2019).

### Limitations and further directions

Whereas the model specified in entropy will be useful in many contexts, we recognize some of its limitations, including ones that pertain generally to inferring ancestry using a `structure`-like model and other forms of model misspecification. Population genetic variation among natural populations arises due to clinal isolation by distance, more abrupt barriers to dispersal that result in actual ‘demic’ substructure within species, or some combination of both (Bradburd *et al.*, 2013; Gompert & Buerkle, 2016). The model in `entropy` and other `structure`-like models do not incorporate clinal isolation by distance and variation and thus, cannot address these issues directly. Linear models for population genetic differences can test for evidence of demic structure beyond what could be predicted from geographic distance alone (e.g., Gompert *et al.*, 2014a; Parchman *et al.*, 2016; Crow *et al.*, 2020). Additionally, alternative models can explicitly parameterize continuous clinal variation and guide understanding of the contribution of demic and clinal variation to population structure (Bradburd *et al.*, 2013, 2016; Battey *et al.*, 2020). When feasible, structured and planned geographic sampling can assist in quantifying the contributions of isolation by distance and demes to population variation. Deviation from the assumed evolutionary model (model misspecification) can be quantified through the correlated differences in a population between predicted and observed genotypes (i.e., correlated residual error), which can guide model choice and interpretation (Garcia-Erill & Albrechtsen, 2019).

This limitation of inferring population structure along a cline may be reduced for mixed-ploidy systems, where we expect some level of genetic differentiation across ploidal levels (even with cross-ploidy gene flow), and `structure`-like models would likely correctly partition individuals of different ploidal levels into different demes. However, clinal variation within a ploidal level in a mixed-ploidy system could still lead to similar problems in inference as specified above.

The *q* model in `entropy` also does not formally include deviations from Hardy-Weinberg equilibrium due to inbreeding (or due to potential double reduction in autopolyploids, see Luo *et al.* 2006 and Bourke *et al.* 2015) and the resulting excess homozygosity of individuals (*F_IS_*) in the prior probabilities for genotype. With sufficient sequencing depth, genotype estimates will strongly reflect the data rather than the prior probabilities, and inbreeding (*F_IS_*) could be estimated in a separate model. Alternatively, the `entropy` model could readily be extended to formally model excess homozygosity. Overall, we are hopeful that the model specification and software implementation might allow others to extend it to additional contexts of interest, including possibly to pools of individuals that are sequenced together, or to chromosomal blocks of ancestry.

Even though the `entropy` model does not explicitly account for allopolyploids, we note that the model can still provide estimates of admixture proportion for loci within separate subgenomes or chromosomes in a higher ploidy individual by treating them as coming from a lower ploidal level. For instance, in allotetraploid individuals with disomic inheritance, we can run a diploid `entropy` analysis on a set of loci coming from one pair of homoeologous chromosomes and a similar analysis on the set of loci coming from the other pair of homoeologous chromosomes, and compare the admixture estimates of the individuals from these two separate analyses as independent realizations of their shared evolutionary history. The pipeline presented in Blischak *et al.* (2017) can be used to obtain `vcf` files with appropriate genotype likelihoods for SNPs in allopolyploid individuals that can then be used as input to `entropy`, by specifying the appropriate ploidal level. The model should probably not be applied to polyploids for which the mode of inheritance is not known, given the potential for spurious clustering due to model misspecification.

The genetic composition of individuals could be the result of a combination of ancient and more recent (i.e., contemporary) hybridization (Gompert *et al.*, 2017; Chaturvedi *et al.*, 2020). Analysis of recent hybridization can benefit from the study of population structure through ancestry-estimation methods such as `entropy`. However, recent hybridization can obfuscate signals of more ancient gene flow (Eriksson & Manica, 2012). Regardless of the extent of contemporary hybridization, alternative models are beneficial to evaluate evidence for more ancient introgression (e.g., Sankararaman *et al.*, 2014; Gompert, 2016; Schumer *et al.*, 2016).

The software for the model is written in C++ using the GNU Scientific Library (Galassi *et al.*, 2009) and the output being written to a Hierarchical Data Format (The HDF5 Group, 2010) file. However, even though the program is written in a low-level language with optimized libraries, given the large size of the estimation problem with typical data sets, the process of converging to a stationary distribution using the Gibbs and Metropolis sampling scheme for MCMC can be time intensive. In future versions of the software, this runtime could be shortened by using techniques like variational inference (as in Raj *et al.*, 2014; Gopalan *et al.*, 2016) and non-negative matrix factorization (as in Engelhardt & Stephens, 2010; Meisner & Albrechtsen, 2018) to arrive at the posterior parameter estimates without using MCMC sampling. However, dealing with the heterogeneity in parameter dimensions that comes with a mixed-ploidy data set will be an algorithmic challenge. For now, in practice we reduce the dimensions of a model run by treating different chromosomes (or other large genome scaffolds) as independent sampling units. This allows one to run separate, parallel analyses of loci on different chromosomes (or scaffolds) that can be distributed across multiple computing cores or nodes, thus enabling us to leverage the high coverage in modern NGS data sets with hundreds of thousands of SNPs.

With these limitations and potential extensions in mind, we find that the `entropy` model can contribute to our understanding of contemporary hybridization and population structure. In particular, the `entropy` model provides a rigorous and beneficial framework for genotype and ancestry estimation from economical, low-depth sequencing data. The model also supports analysis of a wide range of ploidy (from haploid to hexaploid) and mixed-ploidy individuals within a single analysis, which will facilitate a diversity of studies.

## Supporting information

Vignette for entropy software

## Data Accessibility

All simulation and analysis code is available as part of the Bitbucket repository that hosts the source code. The program can be installed via the bioconda channel (https://anaconda.org/bioconda/popgen-entropy) or from source by cloning the Bitbucket repository (https://bitbucket.org/buerklelab/mixedploidy-entropy/), which also houses the on-going developmental code base. A software vignette is part of the Supplementary Material and is also found in the Bitbucket repository. Raw sequence data for *Arabidopsis arenosa* are available at https://www.ncbi.nlm.nih.gov/bioproject/484107.

## Author Contributions

CAB, ZG, EM, DL and TP wrote the diploid model specification and developed the initial software for diploids. The software was tested and improved by DL, PA, EM, TP, and VS. VS extended the model and software to incorporate variable and mixed ploidy, and performed all analyses. VS and CAB wrote the manuscript with input from the co-authors.

## Acknowledgments

We thank colleagues who have contributed to development of the model and its use in various empirical contexts (including C. Nice, J. Fordyce, M. Forister, M. Haselhorst, and S. Lebeis). We thank C. Wagner and K. Hufford for helpful comments on drafts of this manuscript, as well as three anonymous reviewers for providing thoughtful suggestions to improve the manuscript amidst the challenges presented by the COVID-19 pandemic. We thank F. Kolář for sharing the mixed-ploidy *A. arenosa* data from Monnahan *et al.* (2019), and for helpful discussion regarding our reanalysis of their data. This interaction was initiated at the ForBio course “Population Genetics in Polyploids” in 2018, which VS attended with financial support from an NIH INBRE grant to the University of Wyoming. This work was funded in part by the National Science Foundation (DEB-1638602 to CAB). Computing was performed in the Teton Computing Environment at the Advanced Research Computing Center (University of Wyoming, https://doi.org/10.15786/M2FY47).

