## Supplementary material for "Model-based genotype and ancestry estimation for potential hybrids with mixed-ploidy": Vignette for entropy software

### Installing entropy

#### On cluster systems

Similar to most UNIX software packages, you can install `entropy` with the following steps:

1. Download the compressed source code from `https://bitbucket.org/buerklelab/mixedploidy-entropy/src/master/`

2. Unpack the source code with
   `$ tar -zxvf mixedploidy-entropy.tar.gz`

3. If you are on a cluster system, you can load the Intel C++ compiler, the GNU Scientific Library (GSL), HDF5, and AutoTools (both autoconf and automake) as modules
   `$ module load intel gsl hdf5 autotools`

4. Move into the directory with `cd` and install under your home directory in Linux, run
   `$ autoreconf -fi`
   `$ ./configure --prefix=$HOME/entropy`

5. To create the executable, run
   `$ make && make install`

6. The software is correctly installed if you can see the help message displayed when you run the executable from where it is installed
   `$ ./entropy`

#### Using the conda package manager

If you don't already have this package manager, get it installed from here. Most clusters come equipped with `miniconda` modules so you could just load them in to your environment.

1. Add the `bioconda` channel with
   `$ conda config --add channels bioconda`

2. Create a conda environment in which to install `entropy` and other dependencies:
   `$ conda create --name entropy`

3. Install `entropy` with
   `$ conda install popgen-entropy`

4. You can now move in and out of these environments with
   `$ source activate entropy`
   and
   `$ source deactivate`

#### Running `entropy`

The command below runs `entropy` for 20000 steps with 15000 burn-in and stores every 10th step in memory. The input is in the mpgl format (i.e. genotype likelihoods, $m = 1$) and since we are running for mixed-ploidy, we need to include a text file of the ploidy for each individual and locus. We are running for the $K = 4$ model and using initial admixture proportions (for each individual) in the `q4inds.txt` file. We recommend spawning three chains simultaneously (shown for one chain here, with the same length and burn-in period) to assess convergence.

```
./entropy -i auxfiles/final_alfalfa.mpgl -m 1 -n auxfiles/ploidy_inds.txt -k 4
-q auxfiles/q4inds.txt -l 2000 -b 1500 -t 10
-o outfiles/mcmcoutchain1.hdf5
```

#### Command-line options

- -i [file] input data
- -m [0/1] read counts/genotype likelihoods [default=1]
- -n [num/file] single ploidy value/text file with ploidy of loci by individuals [default=2]
- -k [num] putative number of demes [default=2]
- -q [file] file with initial admixture proportions for each individual
- -Q [0/1] intra-/inter-specific ancestry (only available for diploids) [default=0]
- -l [num] number of MCMC steps [default=10000]
- -b [num] number of steps to discard i.e. burn-in [default=1000]
- -t [num] store every [num] step after burn-in [default=1]
- -o [file] .hdf5 output file [default=mcmcout.hdf5]
- -w [0/1] output includes population allele frequencies [default=1]
- -e [num] per-locus error rate if using read counts as input data
- -s [num] scalar for Dirichlet initialization of **q** (inversely proportional to variance) [default=1]
- -p [num $\in$ (0,1)] proposal for ancestral allele frequency [default=0.1]
- -f [num $\in$ (0,1)] proposal for $F_{ST}$ [default=0.01]
- -y [num $\in$ (0,1)] proposal for $\gamma$ [default=0.2]
- -a [num $\in$ (0,1)] proposal for $\alpha$ [default=0.1]

- -r [num] seed for random number generator [default=clock]
- -D [num $\in$ (0,1)] flag to calculate DIC or WAIC estimates [default=0, DIC]

#### Input file format

The software can take either phred-scaled genotype likelihoods or read counts as input. However, we recommend using genotype likelihoods (PL field in the vcf file) since using read counts requires both the counts for each allele but also the per-locus error rate. But this uncertainty is captured by the genotype likelihood (as calculated by a software like GATK, FreeBayes, etc.). To take the data from the vcf and convert it into a readable format by `entropy` (mpgl), we use the the vcfR package (Knaus & Grünwald, 2017). This mpgl file has as the first line, the dimensions of the data set i.e. number of individuals followed by the number of loci. The second line contains a list of the individuals in the data set. The remaining lines start with an identifier for the locus followed by the phred-scaled genotype likelihoods (positive numbers only) for all individuals at that locus. Below is an example for an mpgl file with two diploid (ind01 and ind02) and two tetraploid (ind03 and ind04) individuals with two loci each:

```
4 2
ind01 ind02 ind03 ind04
locus1 0 6 10 0 0 0 0 2 5 10 20 24 6 0 6 24
locus2 0 0 0 10 6 0 24 6 0 6 24 0 0 0 0 0
```

**Note:** The many zero's that are in the mpgl file above, indicate that we have missing information at that locus for that individual. So we assign an equal probability to all the genotype likelihoods (dosages) possible given the ploidy level of the individual. As part of the downloaded package, there is an R script called `inputdataformat.R` that takes as input two vcf files (if you have mixed-ploidy populations) of different ploidy values and combines them to give you one mpgl file that goes in as input to `entropy`. This R script can also be used if you have one vcf file with mixed-ploidy individuals. Two quick checks from the command line to make sure that your input file is correctly formatted are:

- `wc -l infile.mpgl`
  This number should be **one greater** than the number of loci in your data set.
- `awk -F' ' 'NR==3{print NF}' infile.mpgl`
  This number should be **one greater** than the number of genotype classes times the number of individuals in your data set. This should be true for **all** lines in your input data file after line number 3. For example, if you have 180 tetraploids and 60 diploids in your data set, the previous command should give you 1081. Since we have, $180 \times 5 + 60 \times 3 + 1 = 1081$, where 5 is the number of genotype classes for a tetraploid.

This R file also generates initial admixture proportions for $K$ values between 2 and 8 so as to start the chains off in a favorable starting position. These files will be named `qKinds.txt`, where $K \in [2, 8]$. The text files will contain the assignment probability of each individual (in a separate line) to the $K$ different clusters based on the method presented by Jombart *et al.* (2010). To run this script, pass in as arguments the vcf file and the ploidy associated with the file. We recommend splitting up the vcf file based on the ploidy and passing it in as two separate file with a ploidy values associated with each file.

For example, if we have individuals from a mixed-ploidy (2x and 4x) *Arabidopsis arenosa* population that have been split into two separate vcf files:
`Rscript inputdataformat.R arenosa_diploid.vcf 2 arenosa_tetraploid.vcf 4`

Or if you have one vcf file with individuals from both ploidy levels in it:
`Rscript inputdataformat.R final_arenosa.vcf`

**Note:** Ensure that you have the `ploidy_inds.txt` file in the same directory as this R script. Since the R script uses this file to convert the input data from vcf to mpgl format. This R script also requires a lot of memory (i.e. RAM) when working with large vcf files since it needs to read the whole file into memory. So be sure to account for this when running the script.

The next input file that needs to be formatted is the `ploidy_inds.txt` file that indicates the ploidy of each individual (and each locus) to `entropy`. If we have two individuals each from a diploid and tetraploid population (same example as before), the file would look like this:

```
2
2
4
4
```

But there could be cases in which you've sampled loci from the sex chromosomes of the organism and want to model these as haploids instead of diploid. So to let `entropy` know that, you pass it in as a text file that is ind by loci in dimension. So if we have three individuals from a diploid population and five loci, out of which two loci are from the sex chromosomes (which are haploid), then we have:

```
2 2 2 1 1
2 2 2 1 1
2 2 2 1 1
```

**Note:** The most important thing to keep in mind is to ensure that the order of individuals from this text file match the order in the mpgl file.
Similar to the previous check, it is recommended that you run the same checks on your ploidy file before passing it into `entropy`:

- `wc -l ploidy_inds.txt`
  This should be **equal** to the number of individuals in your data set.
- `awk -F' ' 'NR==1{print NF}' ploidy_inds.txt`
  This should be **equal** to the number of loci in your data set.

However, if you do not have mixed ploidy populations in your data set (i.e. all the individuals in your data set are of a single ploidy level), then you can run `entropy` with just one ploidy value in {1, 2, 3, 4, 6}.

### Analyzing output

#### Estimating parameters

The binary `estpost.entropy` contains commands to extract data from the output hdf5 files and present it in human-readable format in text files. The parameter estimates in these text files can then be used in downstream analyses. Below is an example of getting the median genotype estimates with 95% credible intervals across two chains:

```
./estpost.entropy -p gprob -s 0
outfiles/mcmcoutchain1.hdf5 outfiles/mcmcoutchain2.hdf5 -o genoest.txt
```

### Dimensions of parameters

The values for the various parameter (genotype, allele frequency, etc.,) at each step in the chain (after burn-in) are stored in the output HDF5 file.

- genotype (`gprob`): (ploidy+1) x inds x loci
- ancestry (`q`): steps x K x inds
- log-likelihood (`likdata`): steps [DIC] OR steps x (inds x loci) [WAIC]

#### Command-line options

- -o [file] output file [default=postout.txt]
- -p [gprob/q/p/[deviance/likdata]/alpha/gamma/pi/fst] name of parameter to summarize
- -c [num $\in$ (0,1)] credible interval to calculate [default=0.95]
- -s [0/1/2/3/4] type of summary to perform
  - 0 posterior median estimates and credible intervals
  - 1 histogram of posterior samples
  - 2 raw samples from MCMC chains (no summary, just values)
  - 3 calculate DIC or WAIC (based on input file, use -p deviance for DIC and -p likdata for WAIC)

4 MCMC diagnostics (ESS, $\hat{R}$)

- -b [num] number of **additional** MCMC samples to discard [default=0]
- -h [num] number of bins for posterior sample histogram [default=20]
- -v display `estpost.entropy` version number

### Sample workflow

For this section, we'll take a look at how one might run the workflow from beginning to end for a sample data set included under the folder `auxfiles`. This is only a representative dataset meant to capture the mixed-ploidy aspect of the workflow, and hence the final admixture plots are not to be treated as being biologically realistic.

#### Formatting input data

The first step is to format the input data in a way that is readable by the `entropy` program. This is called an mpgl file (specifications provided previously) which basically contains the phred-scaled genotype likelihoods (PL field) for different individuals across all loci in the vcf file (refer to **Note**). Since all the individuals in the `final_alfalfa.vcf` are tetraploid, we can run:

```
Rscript inputdataformat.R final_alfalfa.vcf 4
```

on the command line to generate the `final_alfalfa.mpgl` and the various `qk[2-8]inds.txt` files.

**Note:** Here is where you can filter out loci that are close to each other (in physical distance) to reduce effects of linkage disequiblibrium on the estimates. Based on various analysis, we found that by picking one locus in a window of several kb does the trick. This is a two-way benefit in that we get "truly" independent SNPs in our analysis and we have fewer loci to loop over giving us shorter runtimes. To do this in a practical sense, you can use tools from the R package `vcfR` (Knaus & Grünwald, 2017) to remove loci from the vcf file before converting it to mpgl.

#### Running `entropy`

Now we are ready to run the different configurations of `entropy`. For this, we will need to run the software for K=2 through 8.

```
./entropy -i final_alfalfa.mpgl -n 4 -k 5 -q qk5inds.txt
-l 3000 -b 1000 -t 20 -o alfalfak5chain1.hdf5
```

This is the sample command for K=5, 3000 steps with 1000 burn-in and thinning of 20. The `-n 4` flag indicates that all individuals in the vcf file are tetraploid. This can be replicated for different K values and for different chains since each run is a separate chain (and can be done so simultaneously).

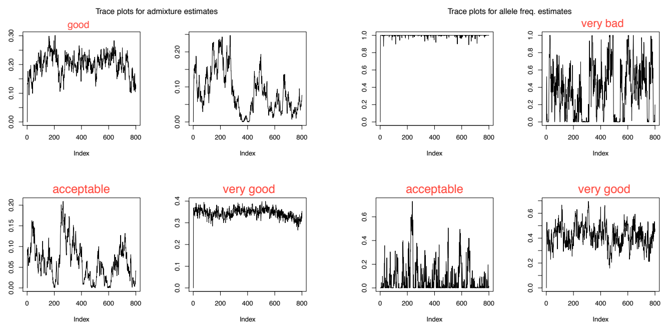


Figure 1: Sample trace plots from an MCMC chain for allele frequency and admixture proportions with annotations to indicate the quality of the chain.

If we have a mixed-ploidy dataset in our vcf file, we need to pass in ploidy values as a text file instead of a single number with the specifications mentioned previously.

#### Assessing convergence

There are two ways we recommend checking for convergence i.e. sampling from a stationary distribution. The first step is to run the chain for as long as you can digest. To get an estimate for how long this may take, run a test chain with the same parameters as above with a shorter length i.e. `-l 1000` instead. This will give you the runtime for a 1000 steps which can be linearly extrapolated with an extra buffer for writing results to disk to 30000 steps by simply multiplying the runtime by 32, for instance. Once the run is completed, you can use the `assessconvergence.R` script to print out trace plots of randomly chosen parameters to see if it resembles a 'good' MCMC chain (needs to look like a fat, hairy caterpillar and not a thin, winding snake). Below is a sample of what you should look for in a good chain.

If most of the trace plots (at least 6 out of 8) look like they are 'good'/'very good' then you can move onto the next step to check for convergence.

To assess convergence of parameters, we can also look at the $\hat{R}$ statistic (Gelman-Rubin statistic/potential scale reduction factor) (Gelman & Rubin, 1992) and see how close it is to 1. This is just a calculation of the variance of estimates within a chain to the variance of estimates across chains. Typically, if the chains have converged to a common distribution $\hat{R} \approx 1$ ($\leq 1.05$). To get an estimate for the $\hat{R}$ statistic for a given parameter, say admixture proportion, we can run:

```
./estpost.entropy -p q -s 4 alfalfak4chain1.hdf5
alfalfak4chain2.hdf5 alfalfak4chain3.hdf5 -o qmcmcdiag.txt
```

This text file also gives us estimates for the effective sample size (ESS) from across our chains. If we have truly uncorrelated samples from our chains, then we can expect this to be close to the number of collected samples.

#### Extracting parameter estimates

Once you have the appropriate $K$ value, it is now just a matter of extracting the parameter estimates from the chains (which are stored in the output hdf5 files). You can do this in two ways: One, the library `rhdf5` (Fischer *et al.*, 2017) can be used in the R computing environment to convert values from hdf5 to R objects. Two, the `estpost.entropy` binary can be used to extract these estimates directly into a text file. The commands for these two options can be found in 1) `hdf52rdata.R` and 2) `hdf52text.sh` under `auxfiles/` of the source repository. The

**Note:** You can make the previously mentioned bash script an executable by using the `chmod u+x hdf52text.sh` command.

#### Obtaining plots of admixture proportions

The admixture/structure plots can be obtained for each value of $K$ in the run using the `plotadmix.R` script provided under the `auxfiles/` folder in the downloaded directory. The script will produce a pdf of the admixture proportions for all individuals and all $K$ values in a stacked barplot fashion. This R script will also output $K$ simple $Q$-matrix text files with individual labels and a single population label file for use with plotting in different softwares like `CLUMPAK` (Kopelman *et al.*, 2015), `pong` (Behr *et al.*, 2016), etc.

#### Running PCA on estimated genotypes

A principal components analysis can be run on the estimated genotypes from the `entropy` model. These genotypes are estimated from an evolutionary process assuming a certain number of parental clusters $K$ with admixture between them. Hence, it provides an alternative approach to estimating genotypes from sequence data from the haplotype-based callers in `GATK` (McKenna *et al.*, 2010), `FreeBayes` (Garrison & Marth, 2012), etc. The estimated genotype values can be extracted using the steps provided in the subsection below, into the R environment directly or into a csv file for raw values (loci $\times$ inds), which can be read into a programming language of your choice.

If all inds in your data set come from the same ploidal level, then the PCA can be run on the resulting genotype matrix directly. However, if there is mixed-ploidy in your data set, it is recommended that the genotype values are scaled by the ploidy (i.e. number of allele copies at a locus) before running the PCA. Basically, this means that the genotype values are divided by the ploidy before running the principal components analysis. This allows us to accurately estimate covariance among inds with different ploidal levels.

#### Choosing the right $K$

The next step in the workflow is to look at which value of $K$ (number of demes) best fits the data. For this, it is recommended that you run `entropy` for all values of $K$ that seem suitable (for example, $K \in \{2, 8\}$). The choice to use the genotype (or admixture proportion) estimates for a certain value of $K$ can be made post-hoc by taking into consideration the caveats pointed out in Lawson *et al.* (2018). This decision can also be aided by the value of WAIC (Watanabe, 2010) that is output by `estpost.entropy` for the $K$ that best fits the data. Below is an example for checking model fit between $K = 2$ and $K = 4$:

```
./estpost.entropy -s 3 -p likdata outfiles/mcmck2chain1.hdf5
./estpost.entropy -s 3 -p likdata outfiles/mcmck4chain1.hdf5
```

(To check whether the **Model WAIC** estimate changes between chains, it is recommended that you run the above command with chain2 and chain3 as well and check the spread of the criterion value.)

There are two components that come together to form the WAIC value: the log-pointwise predictive density and the effective number of parameters. The first term can be thought of the 'deviance' of the model in a Bayesian context. The lower this value, the better the fit to the data. The second term is just the number of effective parameters in the model used to describe the data. As expected, with higher values of $K$ we get better model fit but also an increasing number of effective parameters (which the WAIC penalizes for). So it could be quite possible that `entropy` will best fit the data for a value of $K = 5$ whereas, this number of putative demes could make no sense biologically given your prior knowledge of the system.
