## Supplementary Material for "Model-based genotype and ancestry estimation for potential hybrids with mixed-ploidy"

### Model description

We present a hierarchical Bayesian model to jointly infer genotypes and admixture proportions from sequence data of polyploid and mixed-ploidy populations. This model is implemented in software called `entropy`. This model is similar to the admixture model presented in `structure` (Pritchard et al. 2000, Falush et al. 2003), but with the exception that our model uses genotype likelihood data to incorporate uncertainty arising from sequence data, in the estimation of our downstream parameters. A graphical description of the model is presented in Figure 1 of the main text. The software to sample from the posterior distribution of the parameters (using MCMC) was written in C++, using the GNU Scientific Library (Galassi et al. 2009) and HDF5 (The HDF5 Group 2010). The program can be installed via the bioconda channel (`https://anaconda.org/bioconda/popgen-entropy`) or from source by cloning the Bitbucket repository (`https://bitbucket.org/buerklelab/mixedploidy-entropy/`), which also houses the on-going developmental code base.

Below we provide a more detailed description of the model, with information on the sampling distributions and how it differs from the diploid version in Gompert et al. (2014). We present the two models: the admixture proportion and ancestry complement models, as presented in the main text.

**Model 1 (admixture proportion model)** As described in the main text, the probability of observing the genotype $\mathbf{g}$ is conditional on the unknown population of origin $\mathbf{z}$ of each allele that forms the genotype, and the unknown allele frequencies $\mathbf{p}$ in the source populations, $P(\mathbf{g}|\mathbf{z},\mathbf{p})$. We use genotype likelihoods, $L(\mathbf{g}|\mathbf{X}) \propto P(\mathbf{X}|\mathbf{g})$ rather than raw sequence data $\mathbf{X}$ as model input. The genotype likelihoods are pre-calculated, taking into account the number of reads, number of genotypes, read specific error rate, haplotypic information, etc., given by the sequence data $\mathbf{X}$ (using softwares such as from `GATK`, McKenna et al. 2010, `SAMtools`, Li 2011, or `FreeBayes`, Garrison and Marth 2012). The genotype likelihoods were normalized to sum to 1.

As we restrict our model to work with bi-allelic loci, for an $n$-ploid individual, we can expect to see $n+1$ genotypic states or dosage values at each locus $j$. Each allele at a locus is encoded as either 0 for the reference or 1 for the alternate. The sum at each locus, across all allele copies (e.g., four allele copies in a tetraploid), denotes the genotype at that locus. We can then calculate the probability of each genotype as the product over the probabilities of each allelic state across all alleles, conditional on $\mathbf{z}$ and $\mathbf{p}$. This discrete probability distribution is given as a Bernoulli distribution with a single draw at each allele, repeated $n$ times for each locus in an $n$-ploid individual with probability equal to the allele frequency of the alternative allele in the population of origin $k$.

$$P(g_{ij}|\mathbf{p_j},\mathbf{z_{ij}}) = \prod_{k}\prod_{a=1}^{n} \begin{cases} p_{jk}^{g_{ija}}(1-p_{jk})^{1-g_{ija}} & \text{when } k = z_{ija} \\ 1 & \text{otherwise} \end{cases}$$

Here, $\mathbf{z_{ij}} = [k_1, k_2, \ldots, k_n]$ denotes the local ancestry of the $n$ allele copies for individual $i$, and $z_{ija}$ denotes the local ancestry of the specific allele copy, $a$ in the individual. The term $p_{jk}$ denotes the corresponding allele frequency in the $k^{th}$ source population.

The remainder of our model deviates little from the `structure` *admixture* model with correlated allele frequencies presented in Falush et al. (2003). We specify a set of admixture proportions, denoted by $q_1, q_2, \ldots, q_k$ to indicate the proportion of the individual's genome inherited from each of $k$ source populations. These admixture proportions give the prior for the local ancestry $\mathbf{z}$ in a simple fashion, i.e., $P(z_{ija} = k) = q_{ik}$ for $a \in \{0, 1, \ldots, n\}$. We then place a Dirichlet prior on the admixture proportions for each individual with a scale parameter, $\lambda$, estimated from the data.

The probability of the unobserved allele frequency $p_{jk}$ of locu $j$ in source population $k$ is calculated assuming an $F$-model (as presented in Balding and Nichols (1995)), where the population allele frequency is the result of divergence $F_k$ from an ancestral population, characterized by allele frequency $\pi_j$. We draw $p_{jk}$ from a Beta distribution with shape parameters $\pi_j$ and $(1 - \pi_j)$, both multiplied by $(1/F_k - 1)$. $F_k$ can be seen as a measure of genetic divergence from the ancestral population, analogous to $F_{ST}$:

$$P(p_{jk}|\pi_j, F_k) \sim \text{beta}(\pi_j \frac{1 - F_k}{F_k}, (1 - \pi_j)\frac{1 - F_k}{F_k})$$

The allele frequencies $\pi_j$ are obtained from a symmetrical beta distribution, $P(\pi_j|\alpha) \sim beta(\alpha, \alpha)$. As a result, we obtain a symmetric distribution for the allele frequency in the ancestral population that could take various shapes for different values of $\alpha$, but the distribution is constrained to a mean ancestral allele frequency of 0.5. The hyperparameter $\alpha$ can be seen as a measure of genetic diversity in the ancestral population, and is drawn from a Uniform distribution, $\alpha \sim \text{Uniform}(0, 10, 0000]$. $F_k$ is assigned an uninformative prior $beta(1, 1)$ to indicate equal support for any value of $F_k \in [0, 1]$.

We specify the prior probability for the genome-wide admixture proportion vector $\mathbf{q}$ with a Dirichlet distribution with parameter vector $\gamma = (\gamma_1, \ldots, \gamma_K)$. To assign the same prior probability for each ancestral deme, we specify identical values for all $\gamma_k$. This is appropriate when assuming that neither individuals with ancestry from a single population, nor any particular class of hybrids dominate the hybrid zone. The hyperparameter $\gamma_k$ is drawn from a Uniform distribution between 0 and 10.

Thus, the posterior probability distribution for the `entropy` model is given by

$$P(\mathbf{g}, \mathbf{z}, \mathbf{p}, \mathbf{q}, \pi, \mathbf{F}, \alpha, \gamma|\mathbf{X}) \propto P(\mathbf{X}|\mathbf{g})P(\mathbf{g}|\mathbf{p}, \mathbf{z})P(\mathbf{z}|\mathbf{q})P(\mathbf{q}|\gamma)P(\mathbf{p}|\pi, \mathbf{F})P(\pi|\alpha)P(\phi)$$

where $P(\phi)$ is the joint probability of the terminal parameters in the hierarchy.

**Model 2 (ancestry complement model)** This model is almost identical to the previous model, except in the formulation of admixture proportion. Here, we make use of a matrix $\mathbf{Q}$ instead of the vector $\mathbf{q}$ that is used in `structure`, called the ancestry complement matrix to specify interspecific (or inter-demic) ancestry at a locus, as we shall see below. This model is only available for diploid individuals.

We can obtain additional information on genome-wide admixture by considering a combination of ancestry states at each locus, instead of treating each allele copy as being derived independently from a source population. Therefore, we calculated the probability for locus-specific ancestry jointly for both allele copies (in a diploid) by working with ancestry $\mathbf{z_{ij}}$ as a whole, instead of ancestry for each allele copy $z_{ija}$ separately. The ancestral parameter $\mathbf{z_{ij}}$ can be seen as a $K \times K$ matrix with all its elements set to zero except the element at row

$k = z_{ij1}$ and column $k' = z_{ij2}$ set to one. This means that $\mathbf{z_{ij}}$ is represented as a "one-hot" vector with the index for the corresponding source population denoted by a one, with the remaining entries being zero. This indexing of allele copies to a source population lets us select the corresponding allele frequency for the individual when calculating downstream parameters. The probability of the locus-specific ancestry is then calculated conditional on the genome-wide ancestry complement matrix $\mathbf{Q_i}$ for individual $i$. $\mathbf{Q_i}$ is another $K \times K$ matrix that gives the prior probabilities for genome-wide admixture, or genome composition, for each of the possible states of $\mathbf{z_{ij}}$, with all elements in $\mathbf{Q_i}$ summing to one. The elements on and off the main diagonal give the probabilities for intra-source and inter-source ancestry, respectively. The probability for locus-specific ancestry conditional on genome-wide admixture follows a categorical distribution (or a multinomial distribution with one draw) and is given by

$$P(z_{ij_{kk'}} = 1|\mathbf{Q_i}) = Q_{i_{kk'}}$$

with $k$ and $k'$ giving the rown and column of the $\mathbf{z_i j}$ and the $\mathbf{Q_i}$ matrix. The genome-wide admixture proportion $\mathbf{q}$ is not included as a model parameter but can be calculated marginally from $\mathbf{Q}$ within each interation as

$$q_{i_k} = \frac{1}{2}(\sum_{s=1}^{K} Q_{i_{ks}} + \sum_{t=1}^{K} Q_{i_{tk}})$$

Similar to the previous model, the $\gamma = (\gamma_{11}, \ldots, \gamma_{KK})$ prior is now a matrix instead of a vector, drawn from a Dirichlet distribution with an equal weighting on each ancestral deme.

Thus, the posterior probability distribution for all parameters in this hierarchical Bayesian model is given by

$$P(\mathbf{g}, \mathbf{z}, \mathbf{p}, \mathbf{Q}, \pi, \mathbf{F}, \alpha, \gamma|\mathbf{X}) \propto P(\mathbf{X}|\mathbf{g})P(\mathbf{g}|\mathbf{p}, \mathbf{z})P(\mathbf{z}|\mathbf{Q})P(\mathbf{Q}|\gamma)P(\mathbf{p}|\pi, \mathbf{F})P(\pi|\alpha)P(\phi)$$

where $P(\phi)$ is the joint probability of the terminal parameters in the hierarchy, with the only replacement being $\mathbf{Q}$ for $\mathbf{q}$.

### Calculation of Watanabe-Akaike Information Criterion (WAIC)

The WAIC value is a combination of the log predictive pointwise density (*lppd*), similar to the model likelihood output by `structure`, with a penalization term for the number of parameters in the model (since models with more parameters fit the data better). Consequently, for the IC and the negative log-likelihood, a lower value signifies a better fit. This WAIC value differs from the Deviance Information Criterion (DIC, Spiegelhalter et al. 2002) in that the log-likelihood value is averaged across all posterior samples instead of being calculated on a single average value of the posterior samples. Similarly, the effective number of parameters, which is the penalization term, is also computed using the variance of the log-likelihood (i.e. 'deviance') across all samples. This measure is suggested to work well with a hierarchical model in which the parameters increase in number with the dimensions of the data (Gelman et al. 2014), which is the case in our model.

$$lppd = \sum_{i=1}^{n}(\frac{1}{S}\sum_{s=1}^{S}p(y_i|\theta^S))$$
$$p_{WAIC} = \sum_{i=1}^{n}V^S(\text{log}p(y_i|\theta^S)), \text{where}$$
$$V^S(a_s) = \frac{1}{S-1}\sum_{s=1}^{S}(a_s - \bar{a})^2$$

Then, to get the WAIC value, we calculate $WAIC = lppd - p_{WAIC}$. Here, $p_{WAIC}$ is a measure of the effective number of parameters in the model which is calculated across all data points ($n = loci \times individuals$, is the number of data points i.e., genotypes) and the post burn-in steps in the MCMC chain ($S$, number of draws of posterior distribution). The $V^S$ calculates the variance in our deviance values (i.e., log-likelihood of the data given the parameters).

#### MCMC updates

Below we describe the process for sampling from the posterior for each of our parameters (using various techniques) given the conditional distributions mentioned above. This process also acts as a proxy for the formulation in code with each update step written into a separate function. We will move downward from the graph presented in Figure 1 of the main text. The following text was adapted from the Supplement of Lindtke et al. (2014), with minor changes for dealing with higher ploidal levels.

1. Update $\mathbf{g}$ (sampled from the full distribution)
2. Update $\mathbf{z}$ (sampled from the full distribution)
3. Update $\mathbf{p}$ (Gibbs sampling)
4. Update $\boldsymbol{\pi}$ (Metropolis sampling)
5. Update $\mathbf{F}$ (Metropolis sampling)
6. Update $\alpha$ (Metropolis sampling)
7. Update $\mathbf{q}/\mathbf{Q}$ (Gibbs sampling)
8. Update $\gamma$ (Metropolis sampling)

To implement this sampling procedure, we cycle through each parameter in the model and run the update step for this parameter by holding all other parameters constant at their current value. Once, we are through all the parameters in the model, we will start back up at the ‘top’ of the hierarchy with the likelihood of the sequence data and run through the same sampling process again. The update steps for each parameter are specified in more detail below:

1. Update $\mathbf{g}$:

$$P(g_{ij}|L(g_{ij}|x_{ij}), \mathbf{z_{ij}}, \mathbf{p_j}) = \frac{L(g_{ij}|x_{ij})P(p_{jk}|\mathbf{z_{ij}}, g_{ij})}{\sum_{g_{ij1}=0}^{1} \dots \sum_{g_{ijn}=0}^{1} L(g_{ij}|x_{ij})P(p_{jk}|\mathbf{z_{ij}}, g_{ij})}$$

   Here, $g_{ij} = \{g_{ij1}, \dots, g_{ijn}\}$ and $L(g_{ij}|x_{ij})$ gives the pre-calculated likelihood of each genotype (the input data). For example, in a triploid ($n = 3$), $g_{ij} \in \{000, 001, 010, 011, 100, 101, 110, 111\}$ for each allele copy. $P(p_{jk}|\mathbf{z_{ij}}, g_{ij}) = p_{jk^1}^{g_{ij1}}(1 - p_{jk})^{1-g_{ij1}} \dots p_{jk^n}^{g_{ijn}}(1 - p_{jk^n})^{1-g_{ijn}}$ is the product of the allele frequencies for the first to the $n^{th}$ allele copy in genotype $g_{ij}$ in population $k^1 = z_{ij1}, \dots, k^n = z_{ijn}$, respectively. This update step essentially combines the likelihood of observing a certain genotype given the read data, scaled by the expected frequency of that genotype at that locus (given by the $P(\mathbf{p}|\mathbf{z}, \mathbf{g})$ term).

2. Update $\mathbf{z}$: For the admixture proportion model, we have

$$P(z_{ijk} = 1|g_{ij}, \mathbf{p_j}, \mathbf{q_i}) = \frac{q_i P(p_{jk}|g_{ij})}{\sum_{k=1}^{K} q_i P(p_{jk}|g_{ij})}$$

   where $P(p_{jk}|g_{ij})$ is given in the previous update step (for a certan $z_{ijk} = 1$). Here, we are multiplying two 1-dimensional vectors of length $K$ and dividing each element by the average to obtain normalized values between 0 and 1. This update follows a similar pattern to the update for the genotypes. Here, we sample from a full distribution because we obtain a value for each cluster $k$ in the discrete probability distribution from 1 through $K$. From these vector of values, we perform a single multinomial draw (i.e., $n = 1$) to obtain an index for the putative ancestral cluster.

   Similarly, for the ancestry complement model, we have

$$P(z_{ijk} = 1|g_{ij}, \mathbf{p_j}, \mathbf{Q_i}) = \frac{q_i P(p_{jkk'}|g_{ij})}{\sum_{k=1}^{K} \sum_{k'=1}^{K} Q_i P(p_{jkk'}|g_{ij})}$$

   where $P(p_{jkk'}|g_{ij})$ is given in the previous update step with only two $k$ values since we are dealing with diploid loci and two allele copies, meaning two ancestral populations in $k$ and $k'$.

3. Update $\mathbf{p}$:

$$P(p_{jk}|\mathbf{z_j}, \mathbf{g_j}, F_k, \pi_j) \sim \text{beta}(\pi_j(\frac{1}{F_k} - 1) + r_{ijk1}, (1 - \pi_j)(\frac{1}{F_k} - 1) + r_{ijk0})$$

   where

$$r_{ijk1} = \sum_i \sum_n \begin{cases} g_{ijn} & \text{when } k = z_{ijn}, \\ 0 & \text{when } k \neq z_{ijn} \end{cases}$$

   and

$$r_{ijk0} = \sum_i \sum_n \begin{cases} (1 - g_{ijn}) & \text{when } k = z_{ijn}, \\ 0 & \text{when } k \neq z_{ijn} \end{cases}$$

   give the counts for the alternate and reference allele copies assigned to an ancestral population $k$, respectively.

4. Update $\boldsymbol{\pi}$: Propose a new $\pi'_j$ from

$$\pi'_j|\pi_j \sim \text{Uniform}(\pi_j - 0.1, \pi_j + 1)$$

and accept the proposed value as the new update for $\pi_j$ with probability $\min(1, r)$ if $0 < \pi'_j < 1$, with

$$r = \frac{P(\pi'_j|\alpha)}{P(\pi_j|\alpha)}$$

Using Bayes' rule,

$$r = \frac{P(\alpha|\pi'_j)}{P(\alpha|\pi_j)} \prod_k \frac{P(\pi'_j, \theta_k|p_{jk})}{P(\pi_j, \theta_k|p_{jk})}$$

where

$$P(\pi_j, \theta_k|p_{jk}) = \frac{p_{jk}^{pi_j\theta_k - 1}(1 - p_{jk})^{(1-\pi_j)\theta_k - 1}}{\text{beta}(\pi_j\theta_k, (1 - \pi_j)\theta_k)}$$

and

$$P(\alpha|\pi_j) = \frac{\pi_i^{\alpha-1}(1 - \pi_j)^{\alpha-1}}{\text{beta}(\alpha, \alpha)}$$

with $\theta_k = \frac{1}{F_k} - 1$, and the probabilities for $\pi'_j$ are computed in a similar manner.

5. Update $\mathbf{F}$: Proposal for a new $F'_k$ from

$$F'_k|F_k \sim \text{Uniform}(F_k - 0.01, F_k + 0.01)$$

and accept $F'_k$ as new update for $F_k$ (represented here as $\theta'_k$ and $\theta_k$) with probability $\min(1, r)$ if $0 < F'_k < 1$, with

$$r = \prod_j \frac{P(\pi_j, \theta'_k|p_{jk})}{P(\pi_j, \theta_k|p_{jk})}$$

where $P(\pi_j, \theta_k|p_{jk})$ is given from the previous update step, with $\theta'_k = \frac{1}{F'_k} - 1$.

6. Update $\alpha$: Proposal for a new $\alpha'$ from

$$\alpha'|\alpha \sim \text{Uniform}(\alpha - 20, \alpha + 20)$$

and accept $\alpha'$ as new update for $\alpha$ with probability $\min(1, r)$ if $0 < \alpha' \leq 10000$, with

$$r = \prod_i \frac{P(\alpha'|\pi_j)}{P(\alpha|\pi_j)}$$

where $P(\alpha|\pi_j)$ is given from the previous update step.

7. Update $\mathbf{q}/\mathbf{Q}$: This step involves a simple counting procedure and a single Gibbs update by multiplying a mulitnomial likelihood with a Dirichlet prior, which gives us a Dirichlet distribution with updated parameters.

$$P(\mathbf{q_i}|\mathbf{z_i}, \gamma) \sim \text{Dirichlet}(\gamma_1 + \sum_n \sum_j z_{ijn1}, \ldots, \gamma_K + \sum_n \sum_j z_{ijnK})$$

where $z_{ijn1}$ denotes the local ancestry values (either 0 or 1) for each locus $j$ in an $n$-ploid individual $i$ descended from source population $k = 1$. Similarly, for the ancestry complement model, we have

$$P(\mathbf{Q_i}|\mathbf{z_i}, \gamma) \sim \text{Dirichlet}(\gamma_{11} + \sum_j z_{ij11}, \ldots, \gamma_{KK} + \sum_j z_{ijKK})$$

where we sum over the local ancestry values across all loci $j$ in the genome to obtain an estimate for genome-wide admixture proportion.

8. Update $\gamma$: In the model, all elements of $\gamma_k$ are identical (for both matrix and vector form). We, therefore, propose new $\gamma'$ by proposing a single element $\gamma'_k$ from:

$$\gamma'_k|\gamma_k \sim \text{Uniform}(\gamma_k - 0.05, \gamma_k + 0.05)$$

and accept the new update with probability $\min(1, r)$ if $0 < \gamma'_k \leq 10$, with

$$r = \frac{P(\gamma'_k|\mathbf{q})}{P(\gamma_k|\mathbf{q})}$$

Using Bayes' rule,

$$r = \frac{P(\mathbf{q}|\gamma'_k)P(\gamma'_k|\gamma_k)}{P(\mathbf{q}|\gamma_k)P(\gamma_k|\gamma'_k)}$$

$$r = \prod_i \frac{P(q_i|\gamma'_k)}{P(q_i|\gamma_k)}$$

with the probabilities given in the previous update step. Since we adopt a Metropolis sampling scheme (i.e., symmetric proposal distributions with $P(\gamma'_k|\gamma_k) = P(\gamma_k|\gamma'_k)$), the second component to our update is equal to 1. This allows us to calculate the probability of acceptance, $r$, without considering this second term.

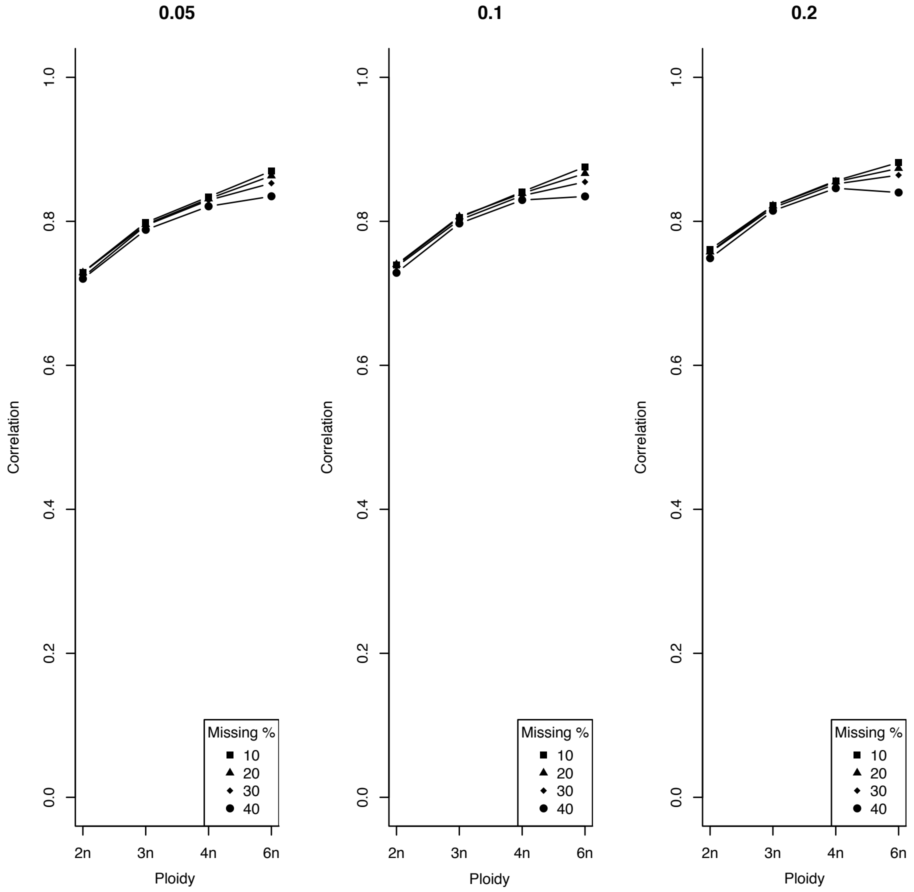


Figure S1: Correlation of estimated **genotypes** at missing sites to true genotypes over varying levels of genetic differentiation $F$ (0.05, 0.1, and 0.2; panes of the plot). There was a slight gain in estimation accuracy when going from 40% missingness to 10% missingness. There was little or no effect of genetic differentiation on estimation accuracy across ploidy levels.

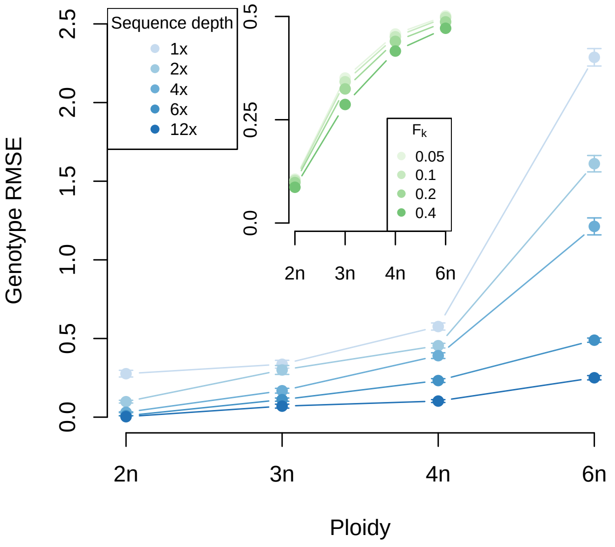


Figure S2: Error in genotype estimation across a range of ploidy decreased with greater sequence depth. The outer plot depicts change in RMSE for different ploidal levels versus sequence depth, across $F = \{0.05, 0.1, 0.2, 0.4\}$ and number of source populations $K = \{1, 2, 3\}$. Error increased with the number of allele copies in polyploids, because the range of variation and possible error in genotype is greater for polyploids than diploids. We found consistently higher error for lower sequence depth (across all ploidal levels). The inner plot depicts the change in RMSE for different ploidy and population differentiation ($F$) across a sequence depth of $n\times$ (with $n$ ploidy and $K = 2$). Error in genotype estimation increased with ploidal level, but was affected very little by the extent of population differentiation.

| Model | WAIC | lppd | neff |
|---|---|---|---|
| K=4 | 3079.1 | -421.5 | 1118.0 |
| K=5 | 3056.1 | -415.9 | 1112.1 |
| K=6 | 3021.5 | -411.8 | 1098.9 |
| K=7 | 2981.9 | -404.4 | 1086.4 |
| K=8 | 2963.0 | -401.0 | 1080.5 |
| K=9 | 2947.3 | -399.4 | 1074.2 |
| K=10 | 2958.5 | -398.1 | 1081.1 |

Table S1: Table of WAIC values for various K models for the *Arabidopsis arenosa* data set with the best-fit model being $K = 9$ (highlighted in yellow). However, Monnahan et al. (2019) found $K = 6$ to be the best-fit, informed by a combination of the Bayesian Information Criterion (BIC, Schwarz et al. 1978) and a similarity index. Similar to other information criteria, the *WAIC* value provides the support for a certain value of K and is a combination of the log-predictive posterior density (*lppd*, similar to deviance in DIC) and the penalization term for the total number of effective parameters in the model (*neff*).

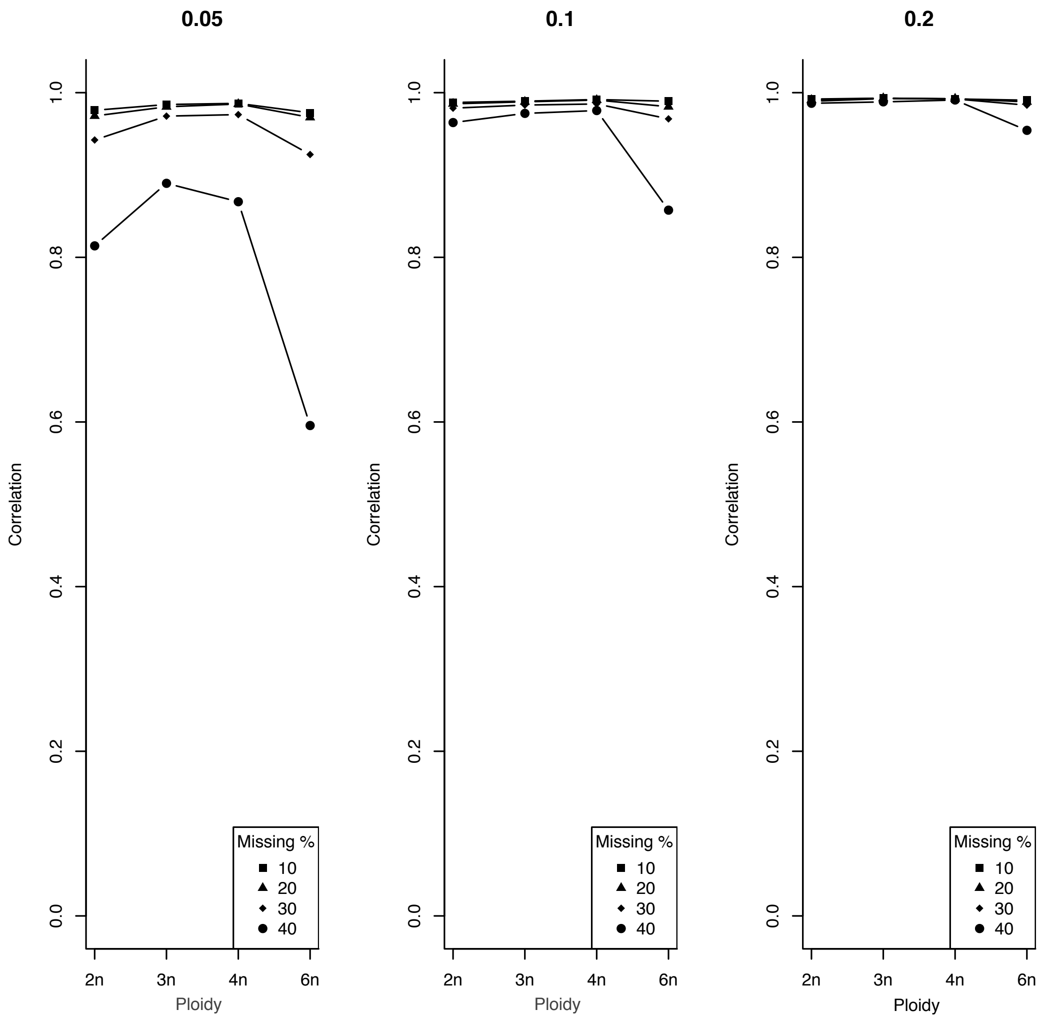


Figure S3: Correlation of estimated **admixture proportion** in individuals over varying levels of missingness and genetic differentiation $F$ (0.05, 0.1, and 0.2; panes of the plot). The percentage of missingness was very important for simulations of demes that were genetically similar ($F = 0.05$) and hexaploid individuals, but even with high levls of missingness, more differentiated parental populations supported highly accurate admixture proportion estimates. The correlation between estimated parameters and the true was very high across ploidy levels (average $\approx 0.97$).

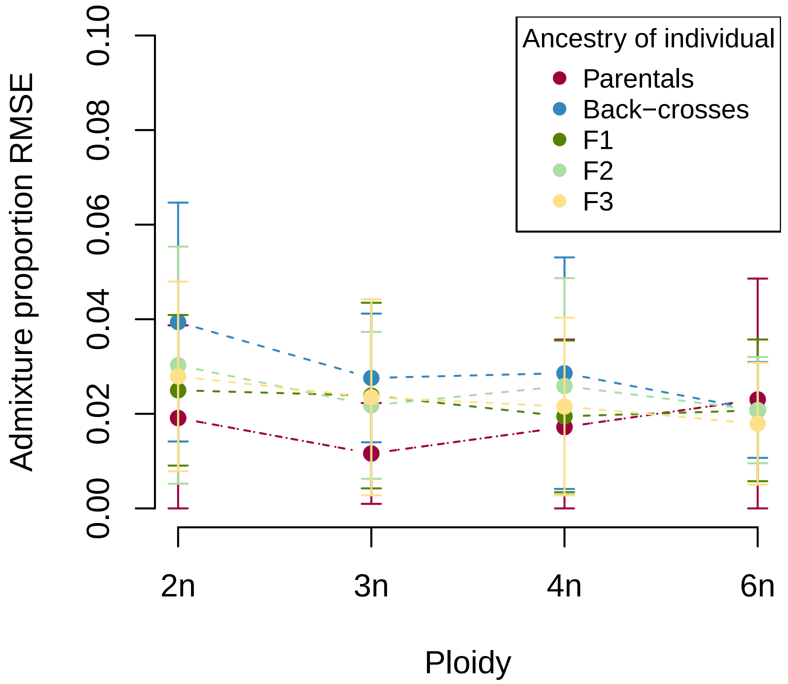


Figure S4: Change in admixture proportion RMSE across ploidal levels. Different ancestry classes in our simulation did not systematically affect the ability to recover true admixture proportions. This is shown for the case of sequence depth equal to $n\times$ (where $n$ is the ploidal level, so sequencing depth equal to ploidy) and across various $F$ values and number of source populations.

| | K=2 & F=0.05 | K=3 & F=0.05 | K=2 & F=0.1 | K=3 & F=0.1 |
|---|---|---|---|---|
| 2n | 0.106 | 0.106 | 0.101 | 0.100 |
| 2n-4n | 0.256 | 0.250 | 0.248 | 0.244 |
| 4n | 0.409 | 0.407 | 0.399 | 0.400 |

Table S2: Error rates for genotype estimates in mixed diploid-tetraploid populations are in between the fully diploid (2n) and fully tetraploid (4n) population. The highlighted row shows that the error for the mixed-ploidy simulation is in between that of the two ploidal levels. This table contains RMSE values for genotypes in diploid (2n), diploid-tetraploid (2n-4n) and tetraploid (4n) populations for different numbers of ancestral demes $K$ and levels of evolutionary divergence $F$.

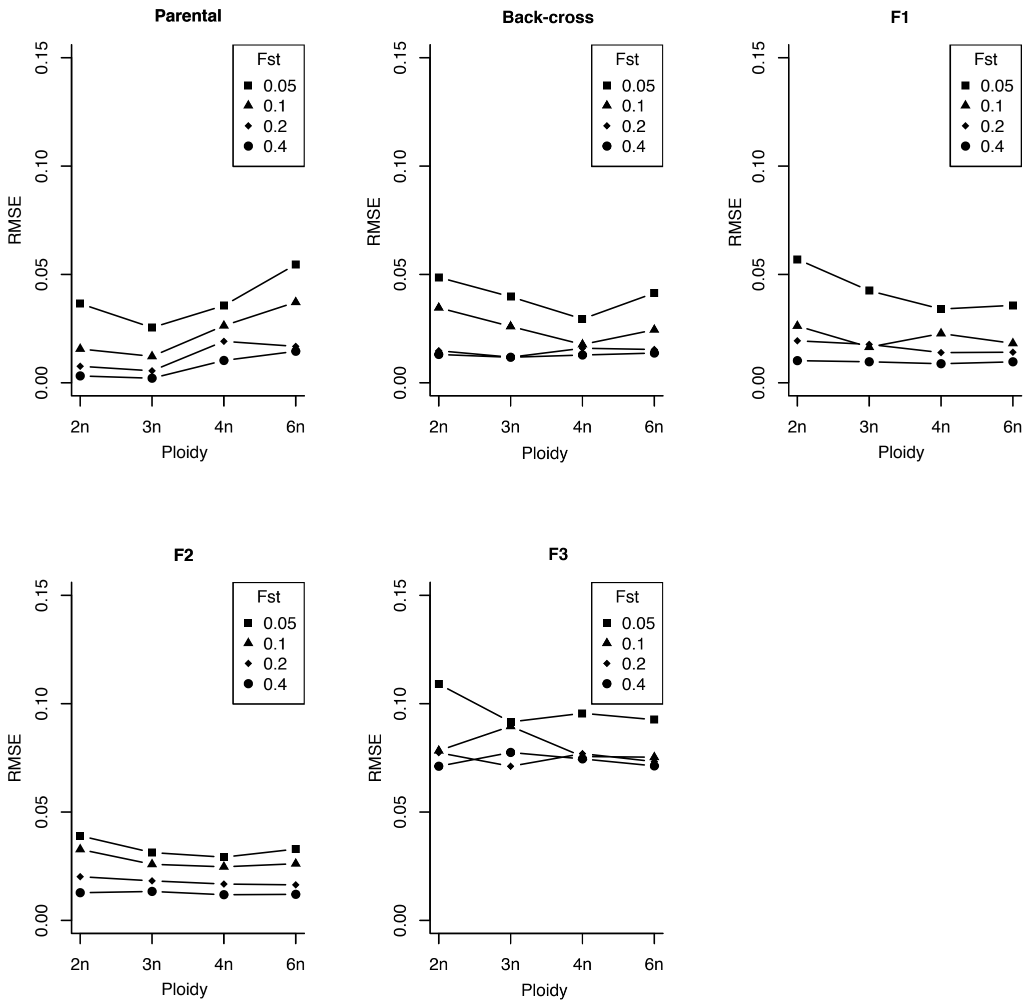


Figure S5: Mean squared error of admixture proportion across five early generation hybrid categories and ploidy levels for varying levels of $F$. Error in estimates decrease with increasing genetic differentiation for all categories of hybrids. The higher overall error with the F3 individuals was because it is harder to estimate the accurate $q$ value given the high realized variance of individual genetic composition around the expectation.

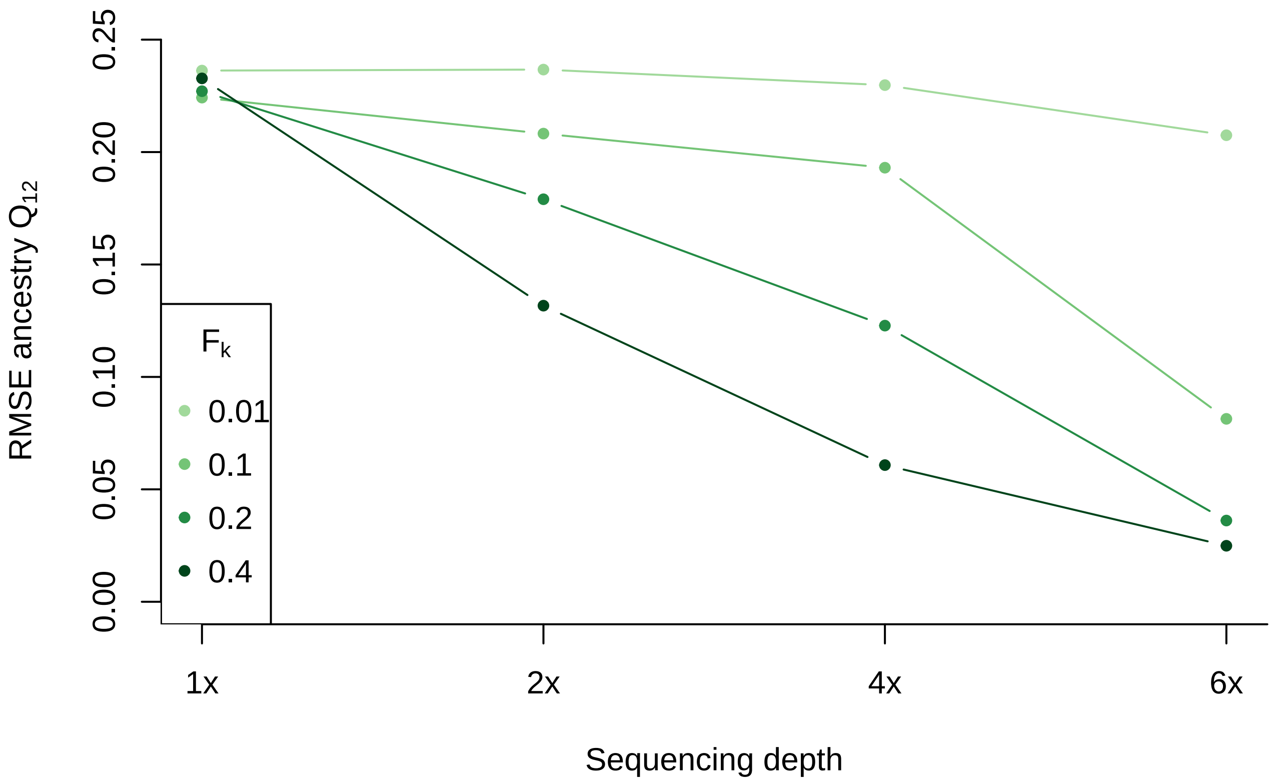


Figure S6: Root-mean squared error for inter-source ancestry $Q_{12}$ (*ancestry complement* model) across a range of sequencing depths for different $F_k$. Error in estimates decrease with increasing sequencing depth for all values of genetic differentiation. Data shown here for 100 simulated diploid individuals for $K = 2$. This matches results from the *admixture proportion* model in this paper and findings from previous empirical studies.

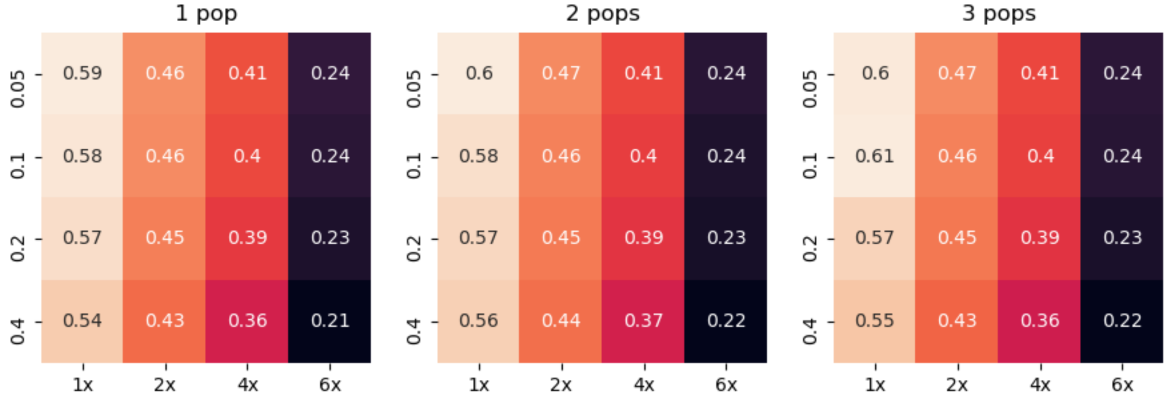


Figure S7: Mean squared error for estimation of genotypes in tetraploid individuals across sequence depths and population differentiation $F$ and number of source populations. The steepest gradient in error was across the sequence depths (i.e., better estimation with greater sequence depth and slight improvement with higher values of genetic differentiation). The lowest error occurred with $F = 0.4$ and $6\times$ sequence depth.

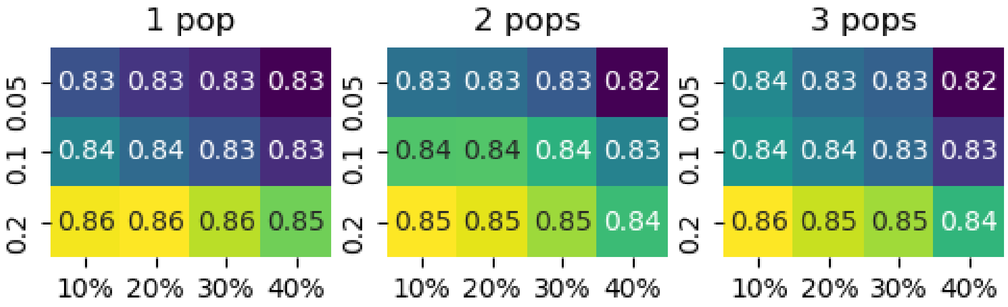


Figure S8: Consistently high correlation of the estimated tetraploid genotypes across degrees of data missingness, three levels of genetic differentiation, and number of source populations. The `entropy` model estimates had a $\approx 83\%$ correlation with the true genotypes at missing sites. Correlations were unaffected or increased slightly with higher differentiation and lower missingness percentage.

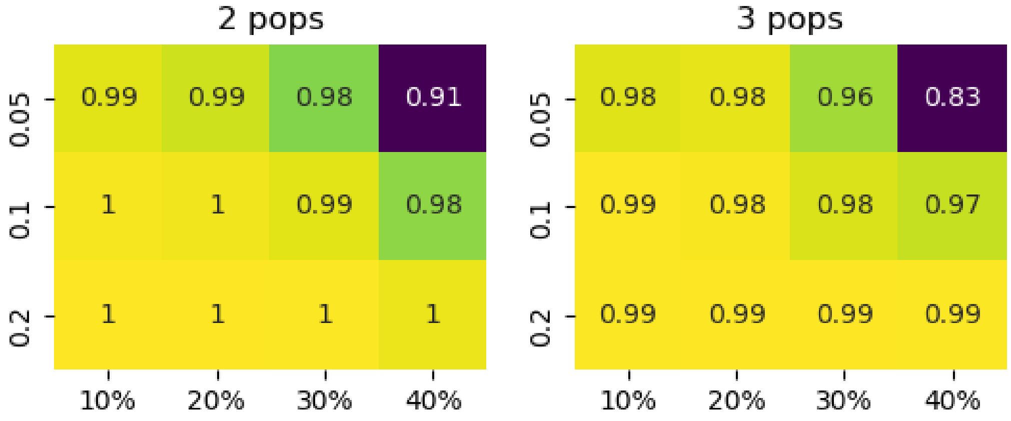


Figure S9: High correlation between estimated and simulated admixture proportion in tetraploid individuals across missingness percentage and genetic differentiation. The model estimated ancestry of individuals with high accuracy, across different levels of missingness in the loci. For example, in a $K = 2$ simulation with $F = 0.05$, the correlation between the true and estimated admixture proportions for individuals with 30% of their sites missing was 0.98. Correlations were lower in simulations with minimal genetic differentiation and high missingness in the data.

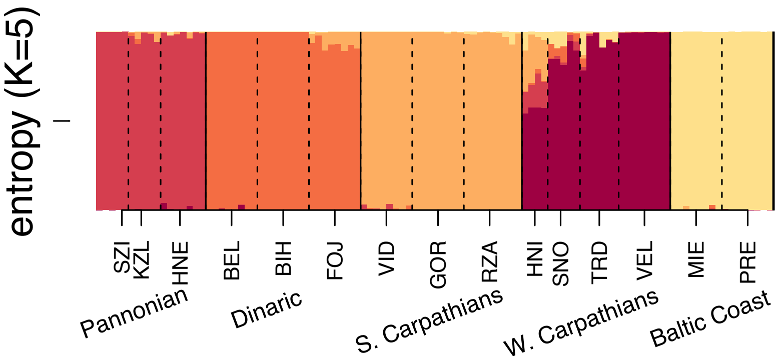


Figure S10: Admixture proportions of only the 105 diploid *A. arenosa* individuals for a K=5 model in `entropy`. The populations in the Pannonian region at the far left (labeled by *SZI, KZL, HNE*) fall into a distinct cluster compared to the rest of the individuals. The Pannonian cluster (red) is genetically the most district from the remaining ancestry groups and was expected to form a distinct group based on the analysis in Monnahan et al. (2019). However, this distinction was not found with the `structure` model applied to the whole data set with a K=6 model (as seen in Figure S11).

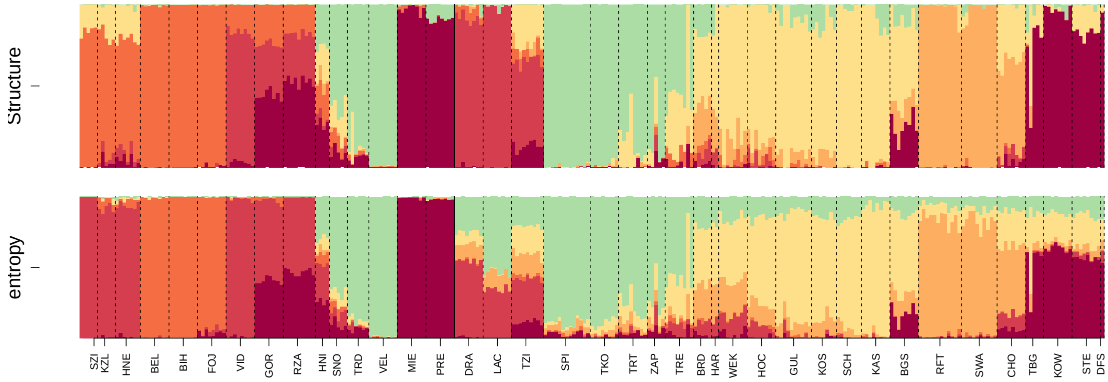


Figure S11: Admixture proportions of all 287 *A. arenosa* individuals for a K=6 model run in `entropy` plotted with population codes instead of regional codes.

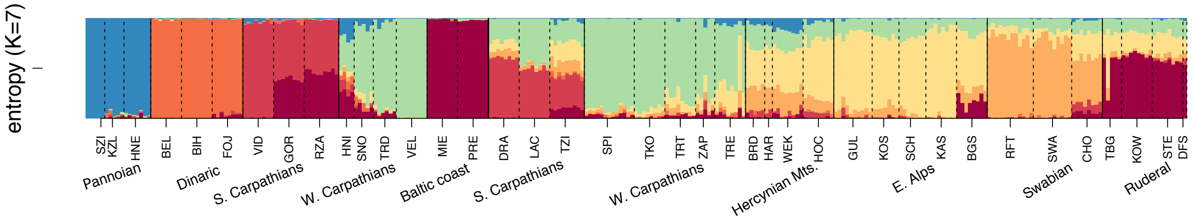


Figure S12: Admixture proportions of all 287 *A. arenosa* individuals for a $K = 7$ model run through `entropy`. The populations in the Pannonian region to the far left (categorized by *SZI, KZL, HNE*) fell into a distinct cluster (blue) compared to the rest of the individuals, further confirming it as genetically differentiated relative to the other populations in the data set.

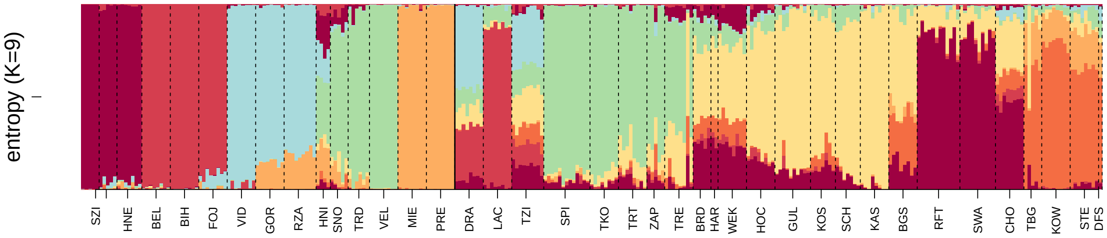


Figure S13: Admixture proportions of all 287 *A. arenosa* individuals for a $K = 9$ model run through `entropy`. The diploid populations in the far left of the plot (colored maroon and red) are characterized as being highly diverged from the rest of the individuals. The presence of multiple source populations in the admixed *DRA, LAC, TZI* tetraploid populations in the Carpathians further emphasize the ability of `entropy` to detect hybrid individuals in a mixed-ploidy zone.

| | | | Assumed | | |
|---|---|---|---|---|---|
| $F$=0.05 | K=1 | K=2 | K=3 | K=4 | K=5 |
| K=2 | 61891.57 | 61321.97 | 61373.16 | 61372.37 | 61430.01 |
| K=3 | 61598.17 | 61179.33 | 60842.83 | 60929.4 | 60931.71 |

| $F$=0.2 | K=1 | K=2 | K=3 | K=4 | K=5 |
|---|---|---|---|---|---|
| K=2 | 61395.55 | 59922.69 | 59892.87 | 59957.28 | 59956.59 |
| K=3 | 62235.11 | 61187.8 | 60205.53 | 60183.08 | 60203.56 |

| $F$=0.4 | K=1 | K=2 | K=3 | K=4 | K=5 |
|---|---|---|---|---|---|
| K=2 | 60957.32 | 53878.83 | 53888.24 | 53867.42 | 53889.24 |
| K=3 | 63773.64 | 58131.46 | 53072.45 | 53083.38 | 53052.75 |

Table S3: The assumed $K$ is found to be equal to the simulated $K$ only 33% of the time for a tetraploid data set. The lowest WAIC values for each of $K = 2$ and $K = 3$ are highlighted in yellow to help for better viewing. This table contains WAIC values to infer best-fit $K$ from `entropy` for different simulation parameters. There is no apparent effect of $F$ on the ability of our model to estimate number of demes $K$. These results differ drastically from the simulated data for hexaploid individuals presented in Table S4.

| | | | Assumed | | |
|---|---|---|---|---|---|
| $F$=0.05 | K=1 | K=2 | K=3 | K=4 | K=5 |
| K=2 | 240331.5 | 240496 | 244859.4 | 244908.7 | 243719.5 |
| K=3 | 241100.1 | 241432.6 | 241246.5 | 242149.9 | 244027.2 |

| $F$=0.2 | K=1 | K=2 | K=3 | K=4 | K=5 |
|---|---|---|---|---|---|
| K=2 | 244342 | 244109.4 | 244231.7 | 247922.2 | 244270.4 |
| K=3 | 243375.4 | 243195.5 | 242924.9 | 243012 | 246588.3 |

| $F$=0.4 | K=1 | K=2 | K=3 | K=4 | K=5 |
|---|---|---|---|---|---|
| K=2 | 264797.5 | 261440 | 261647.6 | 261706.7 | 261645.4 |
| K=3 | 267000.9 | 264881.6 | 262363.8 | 262476.6 | 262560.2 |

Table S4: The assumed $K$ is found to be equal to the simulated $K$ more than 80% of the time for a hexaploid data set. The lowest WAIC values for each of $K = 2$ and $K = 3$ are highlighted in yellow to help for better viewing. This table contains WAIC values to infer best-fit $K$ (for a range) from `entropy` for different simulation parameters, and we see that with higher $F$ we capture 'true' $K$, which differs from the results for the simulated tetraploid data set presented in Table S3.
